# DiPPER2 – a user-friendly pipeline for picking and evaluating taxon-specific PCR primers

**DOI:** 10.1101/2025.06.26.661695

**Authors:** Theresa Wacker, David Studholme

## Abstract

Rapid, specific and sensitive detection is essential for pathogenic microorganisms in contexts ranging from phytosanitary-regulated diseases in agriculture to sepsis management in a clinical setting. Well-established, cost-effective, genomic-sequence-based methods are replacing or complementing laborious, slow and costly culture-based diagnostics. These methods include: end-point PCR and quantitative real-time PCR (qPCR), as well as digital PCR (dPCR) and droplet digital PCR (ddPCR). All these methods require oligonucleotide primers whose nucleotide sequences confer high specificity and sensitivity for the intended target. The most challenging steps in designing these assays include finding genomic sequences that are unique to the target pathogen and absent from off-target organisms, as well as designing primers for those sequences. Here, we present user-friendly, reliable and comprehensive tool DiPPER2 (Diagnostic Primer Picking and Evaluation pipeline for Reproducibility and Reliability), which finds unique diagnostic targets, designs end-point or qPCR primers for them, evaluates the primers *in silico* for specificity and sensitivity, characterizes the diagnostic target and produces a comprehensive and intuitive report. The pipeline has been developed according to FAIR (findable, accessible, interoperable and reproducible) principles and allows for flexible adjustment of primer parameters. It designs primers compatible with qPCR-, dPCR- and ddPCR. DiPPER2 can also evaluate previously designed primer sequences *in silico* for specificity and sensitivity. We use *Pseudomonas amygdali* pv. *morsprunorum* as an example to show that a clade-based, phylogeny-informed approach to diagnostic primer design ensures effective target discovery and specificity.

## Introduction

Rapid and reliable detection of pathogens is paramount in many areas of daily life, including waste- and drinking-water monitoring, testing of regulated seeds, plants and plant products during mandatory phytosanitary controls, and early-stage detection of sepsis in neonates ^1–5^. Compared to slow, costly and labour-intensive culture-dependent methods, culture-independent methods enable cost-effective, rapid and high-throughput detection of pathogens ^1,5^. All sequence-hybridisation based methods, such as PCR, hinge on finding a nucleotide sequence that is unique to the targets (taxa of interest) and the subsequent design of primers that provide sensitivity and specificity for binding to that target sequence.

Before whole-genome sequencing and assemblies became widely available, 16S rRNA and 18S rRNA, as well as ITS (internal transcribed spacer), sequences were frequently used as phylogenetically informative markers for taxonomic identification and classification. However, they do not provide reliable resolution below genus level ^6^. With the steadily increasing number of available whole-genome sequence assemblies, as well as increased computational power, identification of pathogen-specific targets in a reliable and reproducible manner has become less challenging, and several tools have been developed for that purpose. Among those tools are PUPpy, RUCS, Fur in conjunction with Prim and Neighbour (all included in mapro), TOPSI, find_differential_primers (fdp) and SpeciesPrimer ^7–12^.

Both PUPpy and SpeciesPrimer focus on designing primers for annotated genes only, and both were developed for designing primers in bacteria ^8,9^.

Due to using coding sequences (cds) only as an input to identify unique genomic targets, PUPpy relies on accurate annotation ^8^. PUPpy checks target sequences for specificity, however it does not test generated primers for specificity or sensitivity.

SpeciesPrimer includes gene annotation with Prokka into its workflow before using a pangenome approach to identify single-copy core genes for the targets ^9,13^. Like PUPpy, it validates the specificity of the targets, but not of the Primer3 -generated primers, but additionally also includes extensive quality control of the primers it picks ^14^. The use of Prokka restricts its use to prokaryotes (and experimentally viruses) and the annotation and pangenome steps make the overall run very time-consuming ^9,13^.

Unlike both PUPpy and SpeciesPrimer, RUCS uses a k-mer-based approach to find unique genomic regions ^7^. It can also generate primers and tests them. It does so through a combination of using Primer3, BLAST and bespoke filtering steps ^7,14,15^. Unfortunately, RUCS has not been actively maintained since 2022 and lengthy runtimes have been reported previously ^9,16^.

Another tool, TOPSI, runs a series of pairwise alignments of targets to identify regions shared by all targets and conserved regions in targets identified in that manner undergo iterative pair-wise alignments to non-targets to eliminate any regions shared with the non-targets ^12^. After primer-design, specificity is tested using a BLAST search against a database provided by the user. TOPSI requires a Linux cluster with PBS queuing system, and, due to its architecture, the pipeline is slow. Unfortunately, it also is no longer available for download and the webserver, it was hosted on, is only available upon request ^12,17^.

In contrast to the alignment-based approach of TOPSI, the modular fdp pipeline uses an alignment-free approach, which designs primers for all input genome sequences and retains all primers that fall into gene coding regions, excluding primers in all other genomic regions ^11^. They are then cross validated against all targets and neighbours. This step takes a very long time: for the 53 bacterial genomes used in the original paper for primer validation, the cross-validation step took 88h ^11^. A distinct strength of fdp is that it can design sets of discriminatory primers for several subclasses of input sequences simultaneously.

All programs discussed so far encompass all aspects of a primer design pipeline: find unique genomic regions, design primers and test them. This is not the case for Fur in conjunction with Prim, Neighbour and Biobox (as a dependency) ^10^. In this case, Fur is the program used for discovering unique genomic regions and the program Neighbour can be used to find targets and neighbours to feed into Fur. Neighbour relies on the NCBI Taxonomy database to function ^18^. Once Fur has found unique genomic regions in the targets found with the Neighbour program, the program Prim can pick primers and test them for sensitivity and specificity.

Characteristic of all the programs mentioned above is that they have several sequential commands and require substantial user intervention. This can lead to problems with reproducibility, and it limits accessibility. A partial resolution of this and a more user-friendly approach is achieved by the dockerized pipeline mapro, which contains all dependencies and has driver bash script. However, targets and neighbours are still required to be found with the Neighbour program separately and the user needs to generate a relational database. Additionally, only a small subset of the require NCBI *nt* database is included in the Docker container ^19^. The modular and highly customizable structure of the Neighbour, Fur and Prim programs, together with their efficiency, scalability and Fur’s excellent accuracy, can be an asset for the advanced user ^20^. However, unless Docker is used, this makes it far less user-friendly and, in most cases, not easily implementable on a Server (*i.a.* might require conversion to Singularity *etc.*). Additionally, Prim and Neighbour are highly dependent on the NCBI infrastructure.

Here, we present DiPPER2, a fast, user-friendly Python-pipeline which finds genomic regions unique to the target genomes, generates primers, tests the primers for sensitivity and specificity, characterises the unique genomic region found and returns an intuitively comprehensible and comprehensive HTML report that informs the user about the primers it generated (**Figure 1**). Within the DiPPER2 pipeline, Fur is used to find unique genomic regions, leveraging its speed and accuracy ^10,20^. Primers are designed with the widely used and highly customizable tool Primer3 ^14^. *In silico* PCR is performed with the lightweight and fast seqkit amplicon program ^21^. DiPPER2 is available in a testing-only version, which just tests user-provided primers for specificity and sensitivity, and a full pipeline version that also generates unique genomic regions and picks primers.

**Figure 1.**
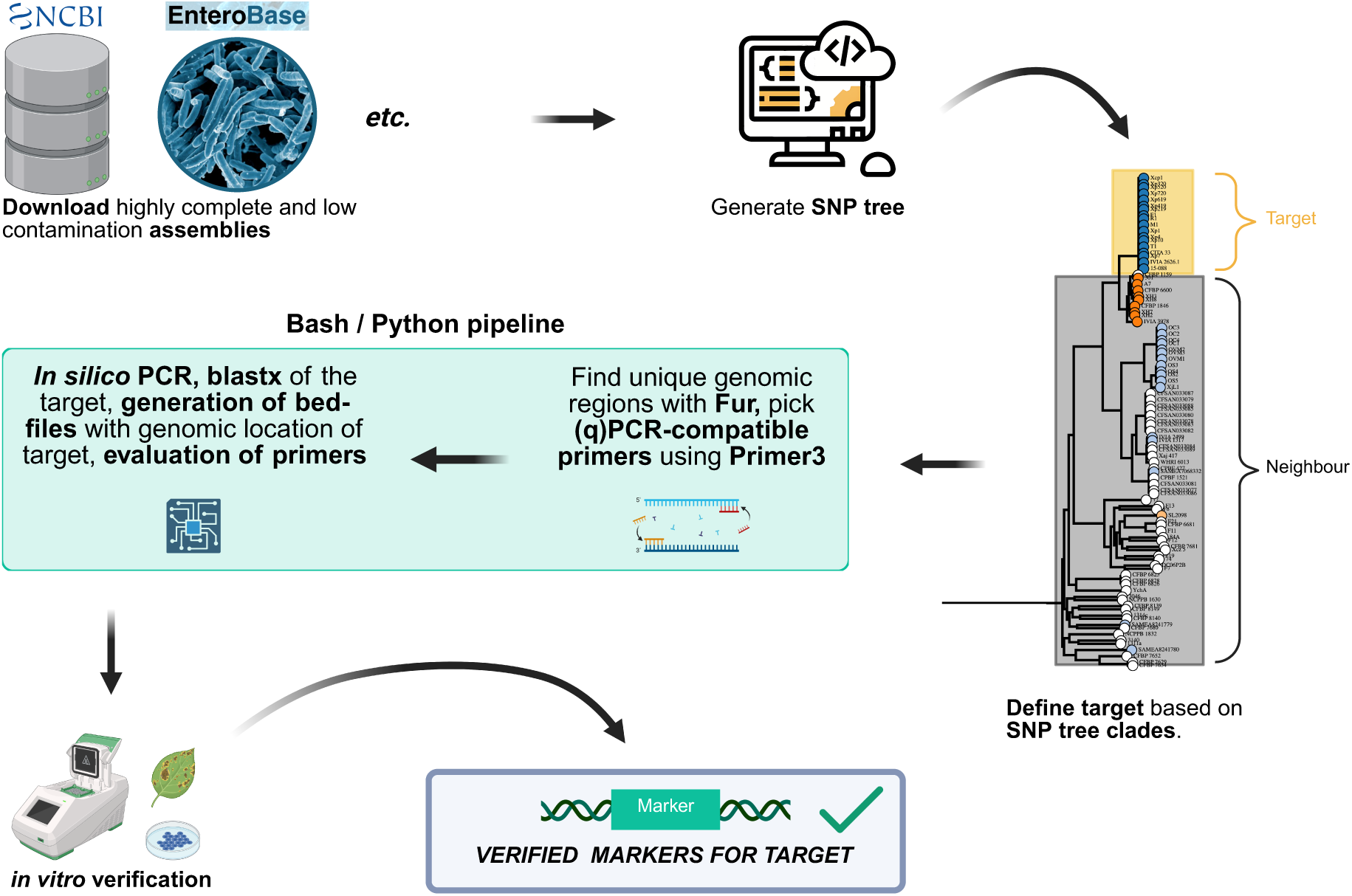
Overview of the full-pipeline mode DiPPER2 workflow.

DiPPER2 is not directly dependent on any specific database and is agnostic to the types of accessions used (*i.e.* does not expect GCA or GCF prefixes), making it more robust to changes in database availability or accessibility.

DiPPER2 is versatile, modularized and user-friendly, and can readily be implemented on a Linux server. As many servers do not run Docker, the containerization method used by *i.a.* RUCS, mapro and SpeciesPrimer, it bridges the gap between a user-friendly and consumer-laptop-compatible and a readily server-implementable configuration and represents a convenient “one-stop-shop” pipeline for the reproducible generation and validation of diagnostic primers, which allows the user to quickly and intuitively find specific and sensitive primers for their target. It takes a phylogeny-informed, clade-based approach and is independent of any annotation, making it more robust to both mis-annotations of CDS and taxonomic misclassifications.

## Results

### DiPPER2 is a fast, user-friendly and modular Python-pipeline

We have developed a modular Python pipeline, that can be run in two different modes: testing-only, to test existing primers *in silico* for specificity and sensitivity, and a full-pipeline mode, which also picks unique genomic regions and designs primers for them, leveraging the highly accurate program Fur for finding the unique genomic regions in assemblies of a target taxon and the versatile, widely used and highly customizable Primer3 for primer picking ^10,14,22^.

DiPPER2’s minimal input for the Bash or Python wrappers that run the pipeline is a list of target genome assemblies that correspond to a clade in a tree and a folder with genome assemblies (both target assemblies and “neighbours”, *i.e.* all other, non-target assemblies). When running the primer-testing mode, a forward and reverse primer are also required. The pipeline can run with minimal user intervention and only requires the user to be able to open a terminal and copy and paste a command. It does not need bespoke data formats and will return standard data formats that are easily useable in other down-stream applications.

Despite its rich functionality, DiPPER2 is not too resource-hungry and is interoperable: DIPPER2 can be run on a laptop with 16Gb of RAM or on a server and works on MacOS, as well as Linux. On Windows it can be run in a virtual machine or in Windows Subsystem for Linux (WSL).

For over 600 assemblies (3.2 Gb of data) DiPPER2 has a reasonably fast runtime (wall-clock time) of under 2h (using max. 6 cores). It provides the user with both a summary of all results in the form of the HTML report as well as a plethora of information, contained in several files and folders (**Figure S1**), including information about the primer characteristics (melting temperature: T_m_ etc.), the unique genomic regions found (including those not used for primer picking), the full Primer3 output, the fasta files containing the primer pairs (default maximum number of primer pairs is 4), *in silico* PCR results, blastx results for the amplicons and BED files with the location of the amplicon in a reference target assembly when blastx results are not available. This way, the user to either refer to the very intuitive, color-coded and comprehensive HTML report or take a deep dive into any step of the pipeline and the information it generated. A customized logger ensures that any errors, warnings and relevant information can be assessed later.

### Rationale of a phylogeny-driven, clade-based approach and reclassification of Pseudomonas amygdali pv. morsprunorum as Pseudomonas avellanae

While not included in the pipeline itself, DiPPER2 highly recommends using a phylogeny-driven, clade-based approach for defining targets and neighbours (**Figure 1**). This avoids the pitfalls of misannotated taxa, which can lead to less or no primers being found or primers being unspecific. If a target is mistakenly defined as a neighbour, unique genomic regions found in the targets now falsely appear to also be found in the neighbours, leading to no or fewer primers being found. Conversely, a neighbour being mislabelled as a target leads to unspecific primers, as the unique genomic region the primers were designed for will find both the targets and the mislabelled neighbour.

For the generation of *Pseudomonas* primers, a SNP tree was generated and revealed that four genome assemblies (GCA_001535755, GCA_002916305, GCA_003698945 and GCA_000145745) annotated as *Pseudomonas amygdali* pv. *morsprunorum* fell within a clade with predominately *P. syringae*, outside of the “main” *P. amygdali* clade, which includes the type strain for this species ( **Figure *2***). Genome assemblies GCA_001535755, GCA_000145745, GCA_003698945 and GCA_002916305 each had an average nucleotide identity (ANI) of more than 97% to *P. avellanae* (**Dataset S1** and **Figure S2**). A threshold of 94-96% is widely accepted to delineate species boundaries, indicating that these accessions are indeed classifiable as *P. avellanae* ^23–25^. Consistent with this, the ANI results indicate that these genomes fall outside of the *P. amygdali* species boundary, sharing between 87.7% and 88.3% ANI with *P. amygdali* (**Dataset S1**). Results of the genome-to-genome distances (dDDH values) are consistent with the ANI calculations and range between 78.1 and 80 in GCA_001535755, GCA_002916305, GCA_003698945 and GCA_000145745 (**Figure** *3*; **Dataset S1** and **S2**). For dDDH, a value of 70 is generally accepted to delineate species boundaries, indicating that the four *P. amygdali* pv. *morsprunorum* should likely be reclassified to *P. avellanae* ^26^.

**Figure 2:**
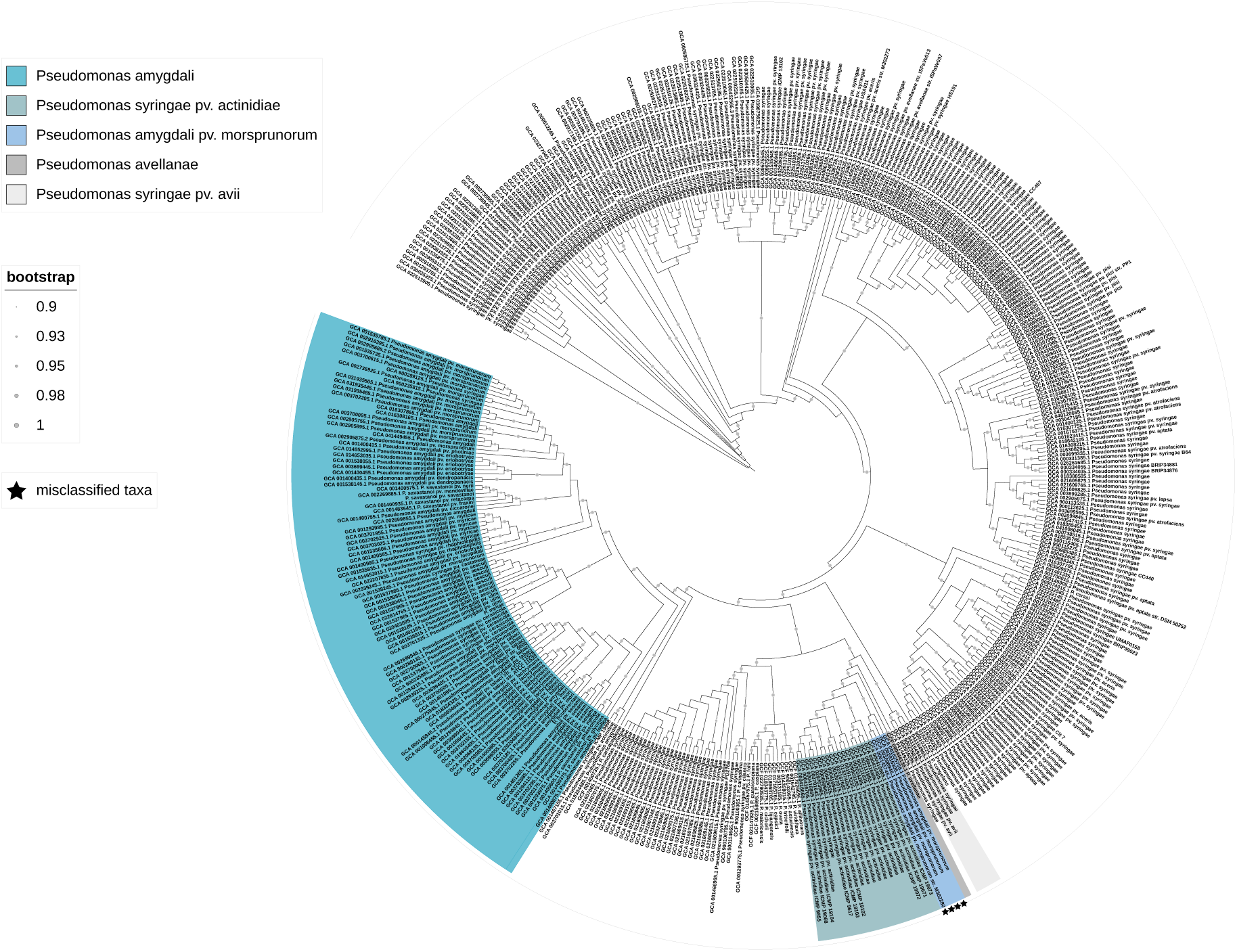
Phylogenetic tree of *Pseudomonas*. SNP tree of *Pseudomonas* assemblies that have a CheckM*^2S^* completeness of ≥90% and a contamination of ≤1%. The 4 mislabelled *Pseudomonas amygdali* pv. *morsprunorum* are marked with black stars.. Tree generated using PhaME^30^ and visualized in iTOL^31^.

**Figure 3.**
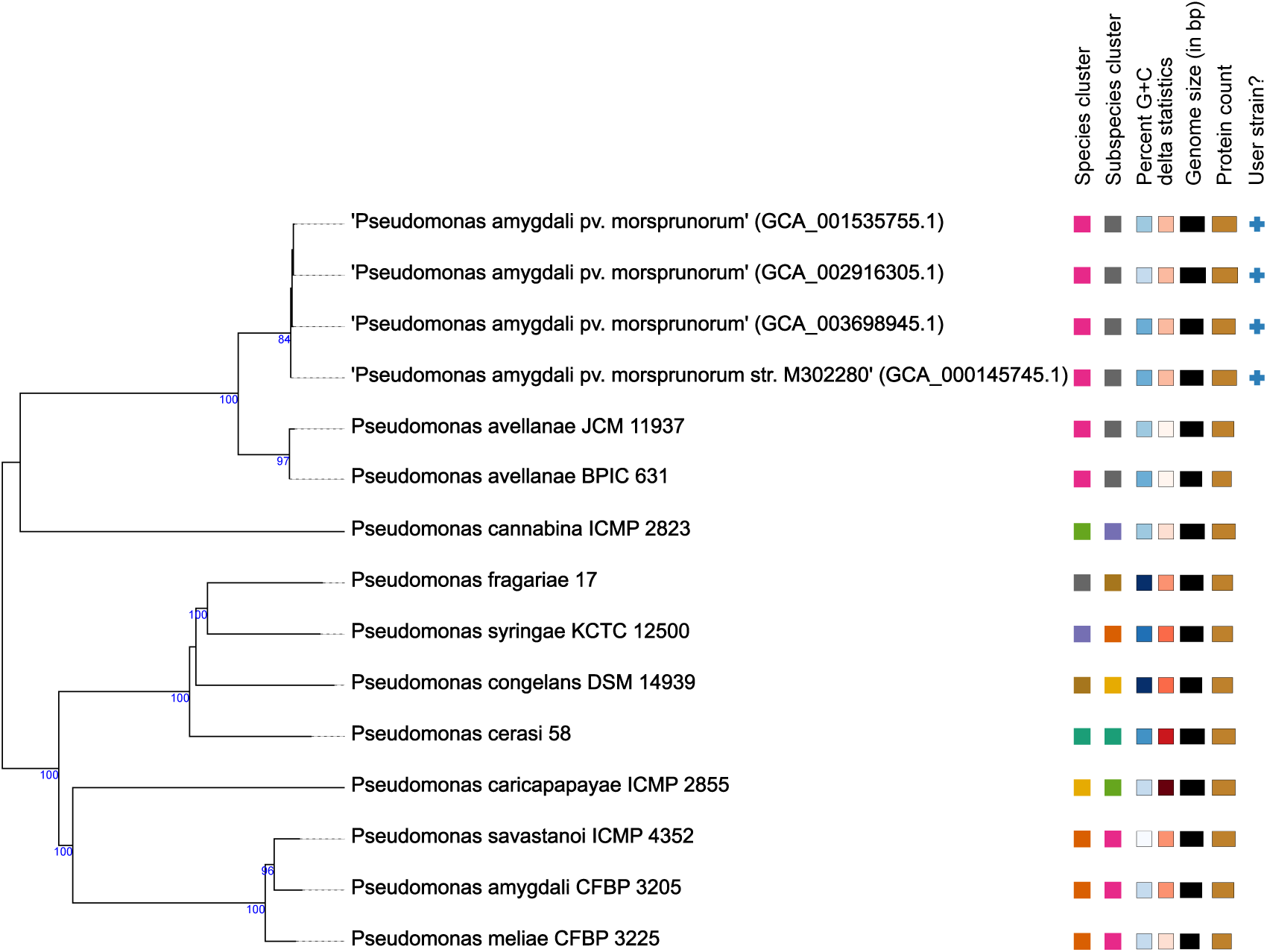
Phylogenetic placement of pv. *morsprunorum* strains in context with type strains of *Pseudomonas* species. The tree was generated by TYGS*^2C^*, which inferred with FastME 2.1.6.1 from GBDP (Genome BLAST Distance Phylogeny) distances calculated from the genome sequences. The branch lengths are scaled in terms of GBDP distance formula d_5_. The numbers above branches are GBDP pseudo-bootstrap support values > 60 % from 100 replications. Mislabelled *P. amygdali* pv. *morsprunorum* are marked with a plus. All genomes apart from the 4 input genomes marked with a plus are assemblies from type material.

### DiPPER2 successfully generates species- and pathovar-specific primers for Xanthomonas *and* Pseudomonas

DiPPER2 was used to generated 6 species-specific and 13 pathovar-specific primer pairs for *Xanthomonas* and *Pseudomonas* (67 qPCR, 36 end-point, excluding primers that failed *in silico* testing; **Table 1**). For each of the taxa listed in **Table 1**, DiPPER2 reported up to 4 primer pairs in an intuitive and comprehensive report (**Figure 4** and **Figure S3**). Of the 109 primer pairs generated, 6 pairs failed specificity or sensitivity testing (0 specificity, 6 sensitivity). All of the failed primers were qPCR primers and failure occurred for 1 or more mismatches in the primer. Of the 8 primer pairs tested *in vitro* for *P. amygdali* pv. *sesami*, 6 were confirmed to be specific and sensitive against a panel of non-target taxa. For 8 primer pairs (4 end-point, 4 qPCR) that were tested *in vitro* for *Xanthomonas euvesicatoria* pv. *vesicatoria,* one end-point primer and 2 qPCR primers were sensitive and specific against a panel of non-target taxa. In this case, the primer pairs that did not produce any amplicons during both qPCR and end-point PCRs failed due to the target amplicon being located on a plasmid that the laboratory strain had lost.

**Figure 4:**
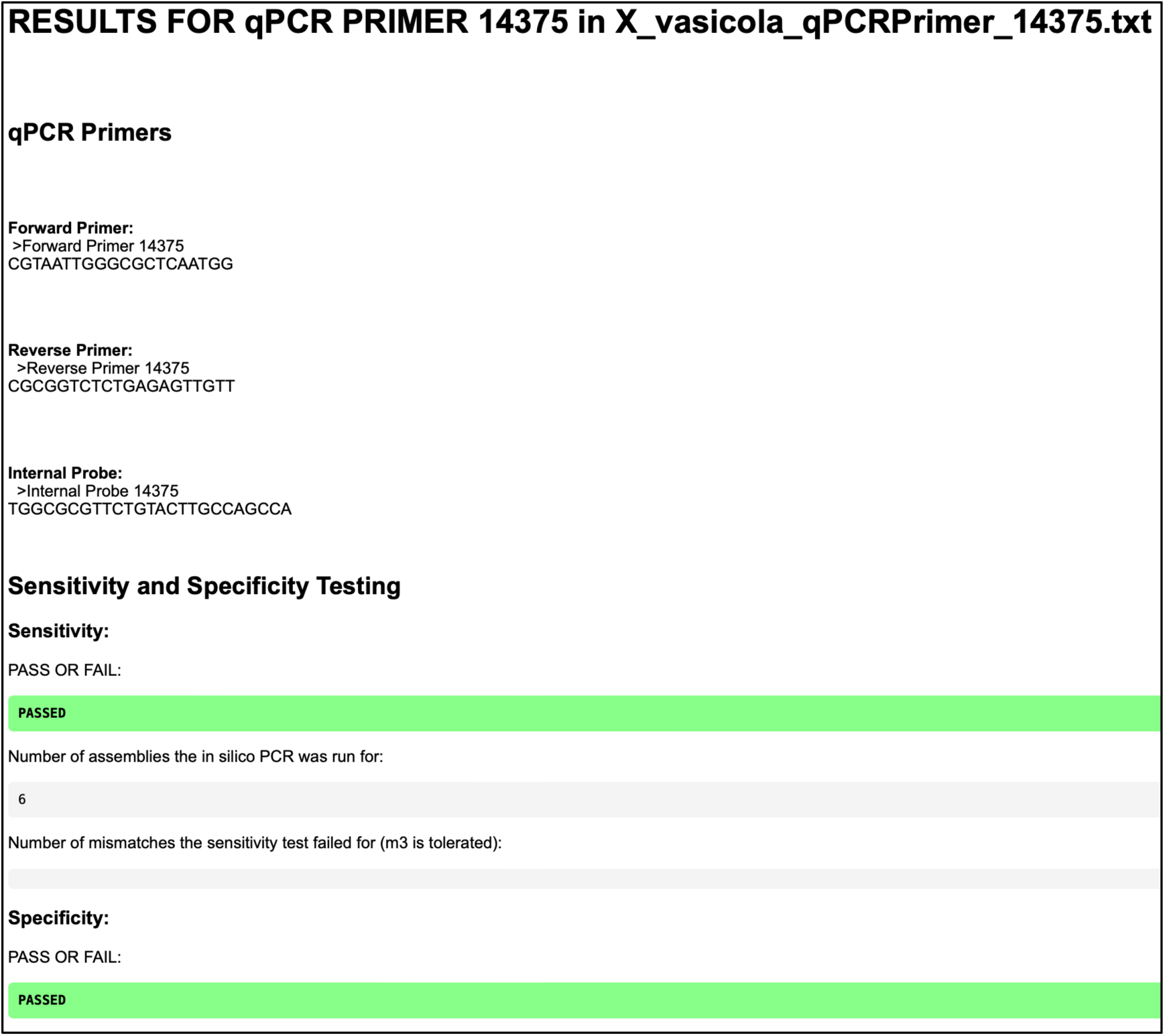
Example of DiPPER2’s Results.html output.

**Table 1:**
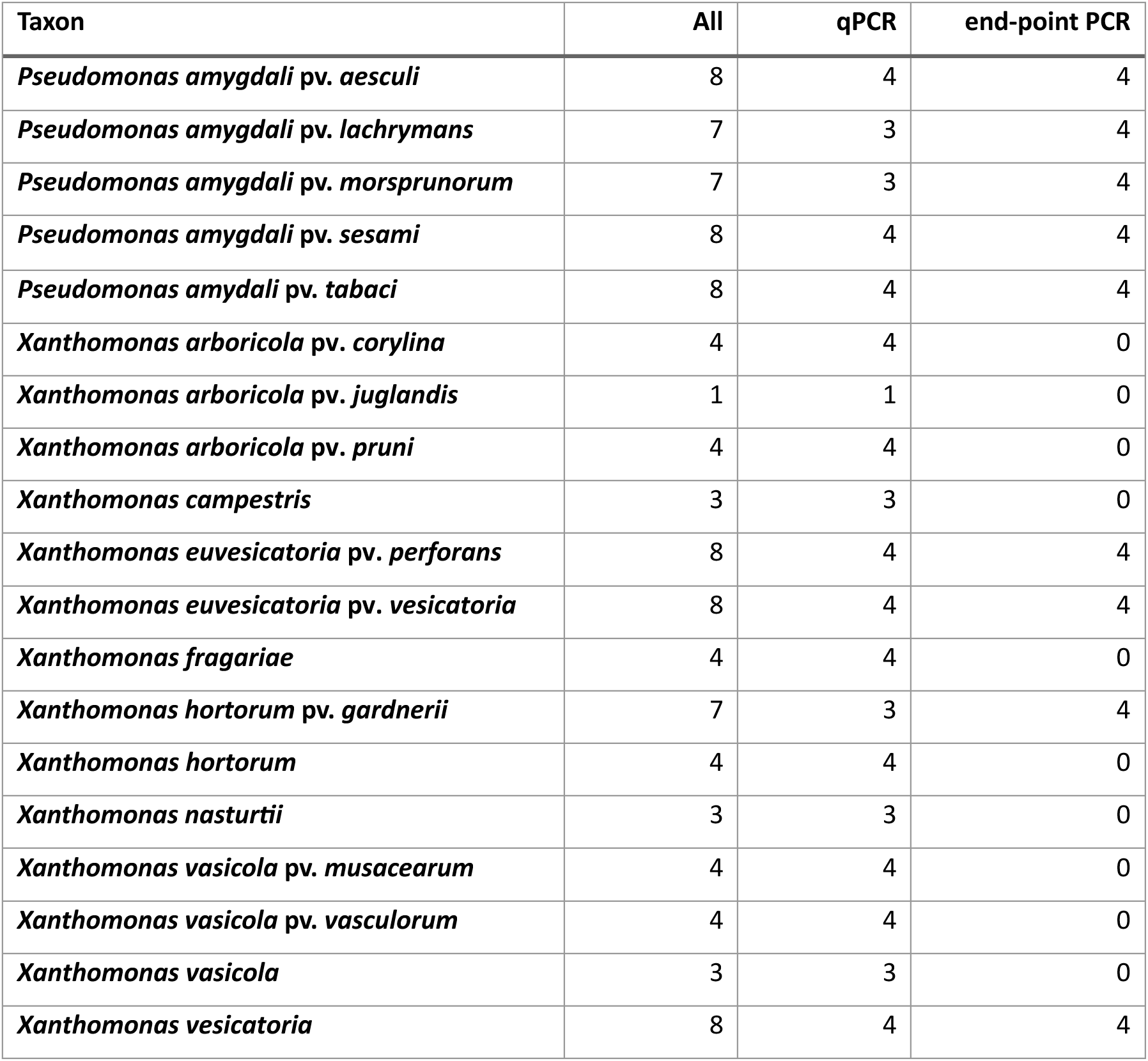
List of primers generated with DiPPER2.

### DiPPER2 can verify specificity and sensitivity of existing primers and ffag potentially problematic primers

Previously published *X. vesicatoria* primers tested with DiPPER2’s testing-only mode primer testing pipeline were specific and sensitive for the panel of *Xanthomonas* tested *in silico* ^27^. However, the previously published *X. vasicola* pv. *vasculorum* primer Xvv3 only passed the *in silico* sensitivity test, but not the specificity test ^28^. When the primer has only one mismatch, it amplifies an amplicon of the correct length in 13 neighbours, 3 of which are *X. campestris* pv. *incanae*, 1 classified as *Xanthomonas campestris* and 9 which are classified as *X. campestris* pv. *campestris* (**Table S1**). The number of correct size amplicons amplified in neighbours is identical for 2 mismatches, for 3 mismatches and above only one correct length amplicon is generated, but numerous long (several kb) amplicons are amplified. This raises doubts about the specificity of this primer.

### Memory requirements and run time analysis

As expected, there is a very strong, positive monotonic correlation (r_s_=1, p<0.001) between the number of assemblies (*i.e.* the amount of data in Gb) and the wall-clock time, as well as between the amount of data and peak RAM usage (**Figure 5**, **FigureS4** and **Table S2**). For over 600 assemblies (3.3Gb of data), maximum runtime was 118min (wall-clock) and the maximum RAM usage was 12Gb. Interestingly, while RAM usage decreased (9.9Gb) for the parallelized version of DiPPER2, runtime increased to 147min (**Table S2**). When makeFurDb was not limited to 6 threads, it required the most RAM to run, as it has a “greedy” configuration and will use as many cores/ threads as it can find. When makeFurDb is limited to 6 threads, seqkit amplicon uses the most RAM.

**Figure 5:**
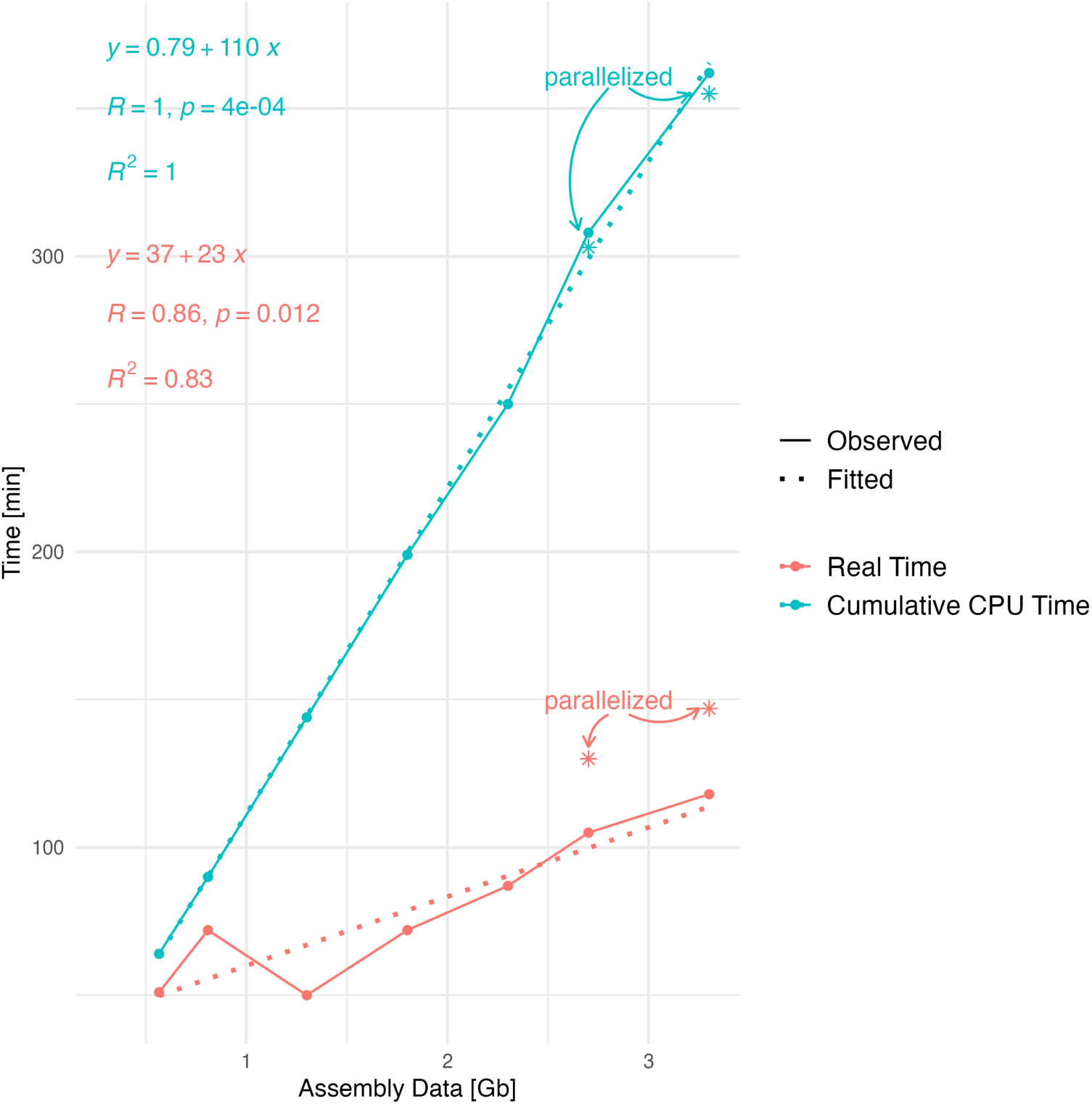
DiPPER2 benchmarking. Real Time (wall-clock time) and cumulative CPU time (additive time of all cores) were measured for DiPPER2 runs with different amounts of assembly data in Gb. 6 target assemblies and between 110 and 660 neighbour assemblies were used as input (0.6-3.3 Gb). Runtimes for the parallelized versions of DiPPER2 were only tested for 2.7 and 3.3 Gb of data and are shown as stars (٭; annotated as “parallelized”).

## Discussion

PCR methods, including qPCR, end-point PCR, dPCR and ddPCR, have revolutionized diagnostics in many areas including plant-pathogen detection, GMO (genetically modified organism) and AMR (antimicrobial resistance) monitoring, medical diagnostics and epidemiological surveillance ^4,5,32–35^. Key to successful PCR applications are specific and sensitive primers tailored to the application.

Here, we present DiPPER2, a fast and user-friendly tool that can generate diagnostic primers for the detection of target species. DiPPER2 can be used to create diagnostic primers that can be used *i.a.* for the detection of plant-pathogens or microbial pathogens with medical relevance, for instance for rapid sepsis screening.

While it does not have a graphical user interface (GUI) “out of the box”, DiPPER2 only requires the user to be able to open a terminal and copy and paste one command. Installation is made as easy as possible with complimentary installation scripts. Once the pipeline runs, it does not require any user intervention. A detailed log of the run allows the user to quickly identify sources of error, should the pipeline fail. It runs on a consumer laptop with reasonably high specifications (requires 16Gb of RAM) and can be readily implemented on a server.

Implementation of DiPPER2 on a phylogenomic platform like PhytoBacExplorer or Enterobase^36^ could further increase ease of use, as this will provide a GUI and embed the tool in a database structure, making downloading assemblies unnecessary and circumventing the necessity of a high-spec consumer laptop with sufficient RAM.

The fact that DiPPER2 is database-agnostic and can be tailored to different databases and accession number systems increases its versatility and ensures that changes in availability and accessibility of databases does not break its functionality. While it does use BLAST to characterise targets, it has a functionality to automatically generate BED files with the coordinates of the target sequence in one of the assemblies of the target microorganism, which allows the user to identify function using a genome viewer like IGV ^37^. While this requires further user intervention and necessitates that the target assembly is annotated, we think that this represents a good fall-back mechanism, should remote BLAST not work and/or the computer or server not be able to communicate with NCBI databases remotely. Due to the modular nature of DiPPER2, this can be changed and expanded, should it become evident that remote BLAST is not the best possible implementation for target characterisation in most cases.

DiPPER2 does not require coding sequences (CDS) files as an input but uses genome assemblies as an input. It does not, like SpeciesPrimer, annotate them itself, increasing its speed and efficiency ^9^. By using genome assemblies rather than predicted proteomes, DiPPER2 avoids biases arising from mis-predictions of protein-coding genes and also allows for finding conserved taxon- or clade-specific regions in the targeted organism’s genome that are not protein-coding, like promotors and 5’-UTRs ^38^. Non-coding regions, especially upstream of essential genes, can be highly conserved. For instance, core promoter elements and transcription factor binding sites are significantly conserved in *Escherichia coli* and were sufficient to recover *E. coli* phylogroups ^39^. Designing primers for non-coding regions can, however, also have drawbacks, either because of ultraconservation that transcends species and genera and renders those regions unspecific or because some non-coding areas can be under relaxed purifying selection ^40^. DiPPER2 specificity testing should exclude ultraconserved regions due to them likely also being in the neighbours of the target clade. With sufficient target assemblies, it is also unlikely that unique genomic sequences found in all target assemblies fall into regions liable to mutation.

Like PUPpy and SpeciesPrimer, which take CDS files or annotate the genome, respectively, DiPPER2 does not discriminate whether a target region is located on a plasmid or chromosome. When bacteria lose a plasmid, this can lead to false negatives, as primers cannot bind and amplify the target region in the organism of interest anymore. Additionally, plasmids can be horizontally transferred. One way to avoid this is to delete plasmid contigs from the assembly FASTA files. However, that is a tedious process. This could be improved by adding a functionality in DiPPER2 that identifies plasmids in the input and excludes them from the search for unique genomic regions in the target.

While DiPPER2 does not automatically generate a phylogeny of the input sequences, it is recommended to be used in a phylogeny-informed, clade-based approach. This is due to taxa being frequently mis-classified in databases and taxonomy changing over time. Changes in taxonomy can result in genome sequences from the same taxon being labelled with different taxonomic names, where those names are synonyms. For example, genome sequences for the plant pathogen *X. vasicola* pv. *vasculorum* are found labelled as *X. campestris* pv. *vasculorum* and *X. campestris* pv. *zeae*. Further confusion can ensue when a taxon comprises heterogeneous organisms. For example, some strains of *X. campestris* pv. *vasculorum* are now considered to belong to *X. vasicola* pv. *vasculorum* while other strains belong to a completely distinct species and are now classified as *X. axonopodis* pv. *vasculorum* ^41^. These scenarios confound attempts to identify taxon-specific genomic signatures. For example, any attempt to identify genomic sequences unique to *X. campestris* pv. *vasculorum* are bound to fail because this taxon encompasses at least two distinct biological entities that are only very distantly related to each other and are more closely related to other taxa than they are to each other within the same taxon.

The pitfalls of going relying on taxonomic information from a database only for inclusion or exclusion of targets and/or neighbours is illustrated by the misclassification of four *Pseudomonas amygdali* pv. *morsprunorum,* which fell into a neighbour clade of *P. amygdali* and were thus excluded from the targets based on phylogeny, albeit otherwise having been included based on the name. Further investigations showed that the four accessions should be reclassified as *P. avellanae.* This highlights that to successfully find unique genomic regions and design taxa-specific primers, a phylogeny-driven approach is advisable and can avoid unspecific primers (due to neighbours being included in the targets accessions) or failure to generate primers (due to targets being included in the neighbour accessions).

DiPPER2 has two modes, one to test existing primers and one to find unique genomic regions, design primers and test those. We show that the testing-only primer testing mode can successfully flag potential issues of existing primers, in this case a primer for *X. vasicola* pv. *vasculorum*, a pathogen causing corn bacterial leaf streak ^28^. The primer was designed using UniqPrimer, which includes optional Primer-BLAST and blastn-based specificity and sensitivity testing, and was experimentally validated *in vitro* ^42^. Using DiPPER2, we show that primers that do not bind perfectly (one mismatch) also amplify the target with a correct size amplicon in *X. campestris*, *X. campestris* pv. *campestris* and *X. campestris* pv. *incanae.* Correct-size amplicons were found in assemblies of which all but one had been deposited in 2023 and were sequenced using PacBio long-read technology (BioProject PRJNA689092). This means that at the time of publication of the primer, cross-validation with Primer-BLAST would not have flagged that problem. One assembly, however, for which *in silico* PCR generated a correct-size amplicon, was deposited both in Genbank and RefSeq in 2013 (GCA_000590335.1). It is unclear why this was not flagged during the Primer-Blast specificity test conducted for the primers in 2017 ^28,43^. The fact that DiPPER2 in testing-only mode was able to successfully flag specificity issues in published primers of a highly-cited paper (cited 51 times; 23/06/2025) highlights the value of *in silico* primer testing: as more and more strains are sequenced and more high-quality assemblies become available, *in silico* testing can flag potential problems, which then can be further investigated and/or tested *in vitro*.

## Conclusion

We present DiPPER2 a fast, user-friendly, modular and versatile Python pipeline, which can either be used to test existing primers for specificity and sensitivity (testing-only mode) or to generate unique genomic regions, pick primers and *in silico* tests them for specificity and sensitivity, as well as assess the target regions of the primers. We show that DiPPER2 can successfully validate published primers for specificity and specificity and report that it has been successfully used to generate 103 primers (end-point and qPCR) for several species and pathovars of *Xanthomonas* and *Pseudomonas*. DiPPER2 is easy to install, can run on consumer laptop with sufficient RAM and is easily implementable on a Server. DiPPER2 is database-agnostic, increasing versatility and maintainability.

Developed with the FAIR principles in mind, it is findable (opensource), accessible (via GitHub and Zenodo), interoperable (MacOS, Linux) and reusable (modular, with GPL-3.0 license, rich metadata and documentation).

## Material and Methods

### Technical Specifications

DiPPER2 was developed and run on Ubuntu 22.04.5 LTS (GNU/Linux 5.15.0-139-generic x86_64). This Ubuntu server was hosted on a Red Hat OpenStack Platform 17.1 server, running an Intel Xeon Icelake processor (2.6GHz) with 32 cores and 64 Gb of RAM. Additionally, it was also developed and run on a consumer laptop with a MacOS Sonoma operating system with 2.4 GHz Quad-Core Intel Core i5 with 4 cores and 16 GB of RAM.

### Software versions

Software and program versions are listed in **Table 2**. Unless otherwise stated, “server” refers to the virtual Ubuntu server, “Mac” to the consumer laptop and “both” when the version is identical on both systems.

**Table 2:** Software Versions.

| Software | Version | Citation/ URL |
| --- | --- | --- |
| **BLAST+** | 2.12.0 (both) | <sup>15</sup> |
| **BowTie2** | 1.3.1 (server) | <sup>44</sup> |
| **CheckM** | 1.2.3(NCBI) | <sup>29</sup> |
| **Conda** | 24.11.3 (server), 23.1.0 (Mac) | <sup>45</sup> |
| **Entrez Direct (EDirect)** | 22.4 (server), 21.2 (Mac) | <sup>46</sup> |
| **FastANI** | 1.32 (server) | <sup>47</sup> |
| **FastTree** | 2.1.11 (server) | <sup>48</sup> |
| **Fur** | 4.1 (server), v4.2-40-gcaafc8f (Mac) | <sup>10,22</sup> |
| **Homebrew** | 4.4.32 (Mac) | https://github.com/Homebrew/brew |
| **iTOL** | 7.2 (web-based) | <sup>31</sup> |
| **iTOL annotation editor** | 1.8 (Mac) | <sup>49</sup> |
| **make** | 4.2.1 (server), 3.8.1 (Mac) | https://www.gnu.org/software/make/ |
| **PhaME** | 1.0.2 (server) | <sup>30</sup> |
| **Phylonium** | 1.6 (server), 1.7 (Mac) | <sup>50</sup> |
| **Primer3** | 2.5 (server), 2.4 (Mac) | <sup>14</sup> |
| **Python** | 3.12.2 (server), 3.9.10 (Mac) | <sup>51</sup>; https://www.python.org |
| **R** | 4.4.1 (Mac) | <sup>52</sup>; https://www.r-project.org |
| **RStudio** | 2025.05.0+496 (Mac) | <sup>53</sup> |
| **Seqkit** | 2.8 (server), 2.6.1 (Mac) | <sup>21</sup> |
| **TYGS** | v400 (web-based) | <sup>26</sup>; https://tygs.dsmz.de |

### Pipeline availability

The pipeline is available on GitHub (https://github.com/ThWacker/DiPPER2) under a GPL-3.0 license. GitHub releases are published on Zenodo (DOI: 10.5281/zenodo.14699017). It is available on the SoftwareHeritage archive with the identifier swh:1:dir:585fd55f9fd6dd486a346fc61376d46d9736157.

### Pipeline architecture

#### Installation

The pipeline can be conveniently installed using the dependency_install.sh script. For MacOS users, Fur will not install automatically and Fur and its dependencies (golang, make, libdivsufsort-dev and phylonium) need to be installed using homebrew as detailed in https://github.com/ThWacker/DiPPER2/blob/main/README.md.

#### Pipeline Input

DiPPER2 assumes that the user has downloaded all genome assemblies that form the pool of targets and neighbours and has generated a tree (or hierarchical clustering) to identify the target clade (or cluster). A text file is required that contains IDs of the target clade assemblies. Together, this represents the minimum input of DiPPER2: a folder with assemblies that are both targets and neighbours in FASTA format (.fa, .fasta, .fna suffix) and a text file with the accessions of the targets. While identification of a target clade based on a phylogenetic tree and/or clustering is not strictly necessary, we highly recommend this phylogeny-driven approach for best results. Mis-annotated neighbours that fall into the target taxon’s clade will result in substantially fewer found unique genomic regions that can be used for the primer design.

Should the user want to test existing primers not generated by DiPPER2, those need to be available as input strings as well.

#### Pipeline workflow

Once these pre-requisites are met, the user has the option to run DiPPER2 as Python or Bash wrappers in two modes:

1. **Testing-only mode:** the DiPPER2 pipeline will test the user-provided primers (forward and reverse as strings) for sensitivity and specificity and characterise the target amplicon. A HTML and a text report are generated.
2. **Full pipeline mode:** the DiPPER2 pipeline will find unique genomic regions that are present in the targets but not the neighbours, design primers, pick the four best-scoring primer pairs, test them for specificity and sensitivity *in silico*, characterize the target sequence that is amplified by the primers and generate a HTML and a text report.

Once the pipeline is started, the wrapper module will establish whether the dipper2 Python environment is already present; if not, it will create the dipper2 environment. It activates the environment, establishes the Python path and version, checks whether all mandatory arguments are provided and starts the first module (**Figure 6A**). This is the same for both modes.

**Figure 6:**
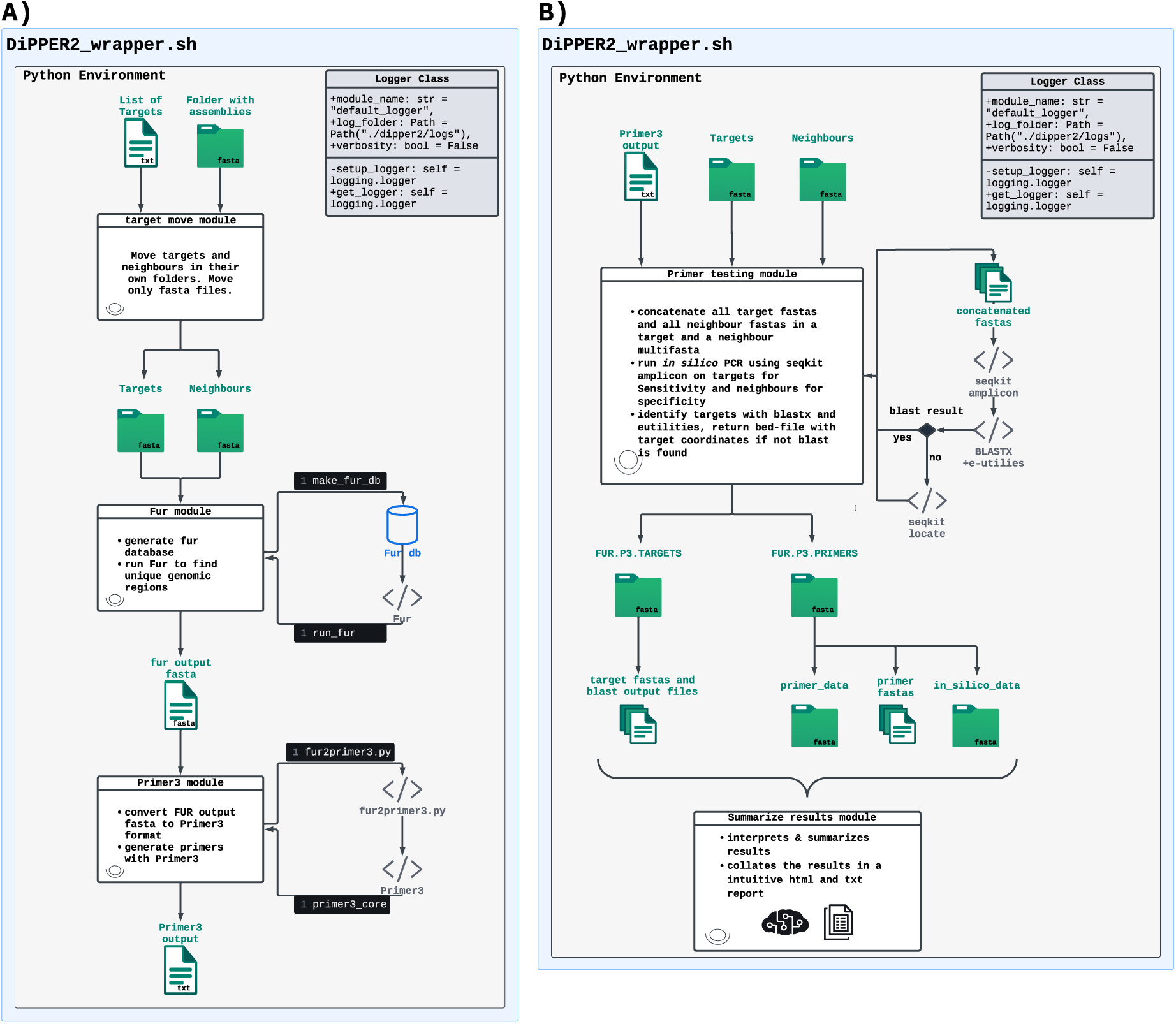
DiPPER2 architecture and workflow. A) Finding of unique genomic regions and primer picking as part of the full pipeline mode B) Primer testing and target characterisation. This is the entry point for the standalone mode and the second part of the full pipeline mode run.

The first module sorts the targets and neighbours found in the user-provided folder with the input assemblies in two folders: FUR.target and FUR.neighbour. This module is called in both modes.

For the full pipeline mode wrapper (DiPPER2_wrapper.sh or.py), the pipeline continues with calling the second module which finds unique genomic regions to design primers on. It leverages efficiency, scalability and accuracy of the Fur program and uses makeFurDb and fur to find unique genomic regions ^10,22^. Fur outputs the consensus sequences of the unique genomic regions in a FASTA-formatted output file (with the suffix _FUR.db.out.txt). It also provides the user with some statistics on how many sequences it finds in each of its three computational steps (Subtraction_1, Intersection, Subtraction_2)^22^; this can be found in the FUR_Summary_output.txt file (**Figure 6A**).

Next, the wrapper calls the third module, the primer design module, in full pipeline mode. This module designs the primers using Primer3. As Primer3 requires a bespoke input format and cannot take as input the the FASTA-formatted output file of Fur, the module calls fur2primer3.py, which converts Fur’s output into a Primer3-compatible one. If the user has supplied primer parameters (for instance requesting changes to the default values in **Table 3**), and if the user has requested qPCR primers (-q flag, default: no (n)), these will also be used by fur2primer3.py to configure Primer3 ^14^. The default values for Primer3 flags that parameterize the primer design with Primer3 are listed in **Table 3**. The primer design module subsequently calls Primer3 and the run of that program produces a bespoke output which is converted by the primer design module in a more interoperable output, including extracting primers, targets and primer characteristics from it. For each picked pair of primers, Primer3 reports a penalty score called PRIMER_PAIR_0_PENALTY. The score is calculated as the weighted sum of normalized deviations of a primer’s properties (melting temperature, G+C content, product size etc.) from the specified optima, plus thermodynamic penalties for hairpin, self-dimer, and cross-dimer formation. It favours thermodynamically stable primers, and lower scores represent better primer characteristics. DiPPER2 selects the 4 lowest scoring primer-pairs for subsequent specificity and sensitivity testing. Primers that are not selected can still be manually reviewed and are found in the file with the suffix_FUR.db.out.primers.primer3_out.txt. Primers pairs are saved in their own FASTA-formatted files and put into the FUR.P3.PRIMERS folder. If qPCR primers were requested, these FASTA-formatted files also contain the internal probe. The characteristics of each primer (Tm etc.) are deposited in files having the same unique identifier as the primer FASTA files in a folder called primer_data. Target sequences that represent the unique genomic regions the primers were designed for and their unique identifiers are saved in the folder FUR.P3.TARGETS.

**Table 3:**
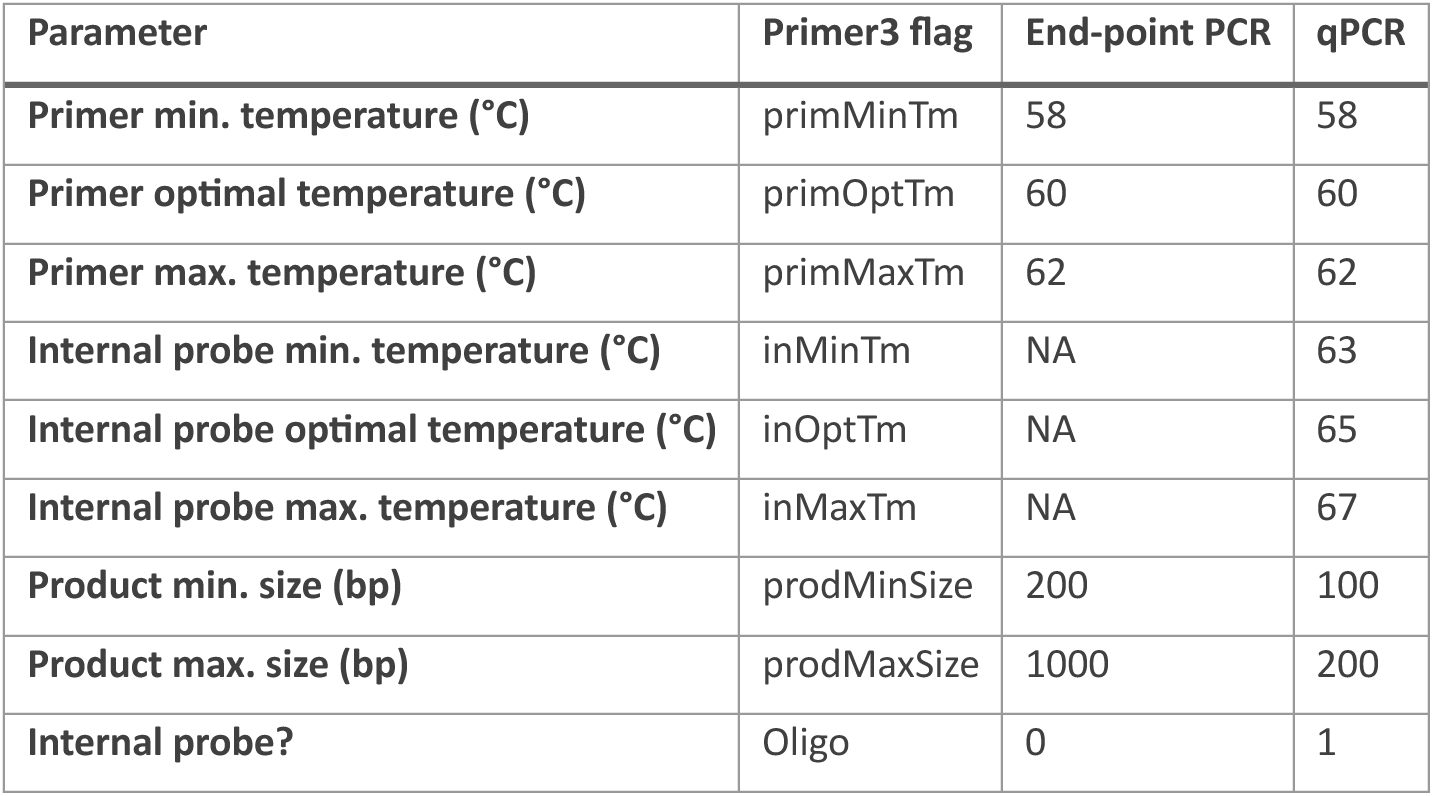
Primer3 default parameters 0=no and 1=yes for the Oligo flag.

| Parameter | Primer3 flag | End-point PCR | qPCR |
| --- | --- | --- | --- |
| Primer min. temperature (°C) | primMinTm | 58 | 58 |
| Primer optimal temperature (°C) | primOptTm | 60 | 60 |
| Primer max. temperature (°C) | primMaxTm | 62 | 62 |
| Internal probe min. temperature (°C) | inMinTm | NA | 63 |
| Internal probe optimal temperature (°C) | inOptTm | NA | 65 |
| Internal probe max. temperature (°C) | inMaxTm | NA | 67 |
| Product min. size (bp) | prodMinSize | 200 | 100 |
| Product max. size (bp) | prodMaxSize | 1000 | 200 |
| Internal probe? | Oligo | 0 | 1 |

After the successful completion of the third module, the wrapper calls the primer testing module (**Figure 6B**). This is also the entry point of the testing-only mode, which can be run using the DiPPER2_standalone_primer_test.sh or DiPPER2_standalone_primer_test.py wrapper. The primer testing module performs specificity and sensitivity tests for each primer using seqkit amplicon from the seqkit program ^21^. As primers do not necessarily bind perfectly *in vitro* (*i.e.*they bind despite not exactly complementing the target sequence), the sensitivity tests are iteratively run with tolerances of 0-3 mismatches in matching primers, while the specificity tests are run with allowing 0-4 mismatches.

Primer pair targets are then characterized using blastx ^15^. If blastx does not return significant mathces, seqkit locate is used to find the genomic coordinates of the target, using either a user-provided reference or the longest target assembly. The resulting BED file can be assessed by the user via a genome viewer such as IGV (Integrative Genomics Viewer)^37^.

The information generated by the modules is then bundled and interpreted by the last module, which produces a comprehensive and intuitively understood HTML report (**Figure S3** and **Figure 5**). The module considers a primer to have passed sensitivity testing when the right size amplicon is amplified in every target. It tolerates wrong sized amplicons for *in silico* PCRs with 3 mismatches allowed.

Specificity tests are only considered as “passed” when there are no correctly-sized amplicons generated in any of the neighbour genomes. Amplicons of the right size in any of the neighbour genomes are not tolerated, regardless of the number of mismatches allowed in *in-silico* primer binding. For targets that produced blastx results, the accession of the BLAST hit with the highest bit score is returned and its annotation is retrieved using efetch from Edirect ^46^. A direct link to the NCBI database entry, if applicable, is also provided in the report.

### Parallelization of primer testing module

The parallelized version of DiPPER2 (DiPPER2_wrapper_parallel.sh) does not concatenate all targets and neighbour assembly FASTA files and run the *in silico* PCR on those but uses the multiprocessing package to run seqkit amplicon on individual assemblies in parallel. To make sure this does not overwhelm the computer or server, the Primer_testing_parallelize.py script checks whether sufficient memory is available before spawning each process. Additionally, the script monitors memory consumption of each of the workers and will kill the process if it exceeds a defined limit.

### Resource consumption

Resource consumption was measured for datasets consisting of between 116 and 666 assemblies (6 *Xanthomonas vasicola* target genomes plus 660 neighbours, chosen as all assemblies with ≥90% CheckM completeness and ≤1% contamination; 0.6-3.3 Gb; **Dataset S3**) on the Ubuntu server (specifications as described in technical specifications) with all programs using default number of threads, other than makeFurDb, which was limited to 6 threads. Wall-clock time and cumulative CPU time were measured using the time command. RAM usage was measured using a bespoke Bash script (memory_monitor.sh, that can be found on the accompanying https://github.com/ThWacker/DiPPER2_publication repository) which uses ps to get the resident set size (rss). Results were visualized in R using the profiling_visualization.R script found in the same repository.

### Verification of specificity and sensitivity of published primers

As an example of DiPPER2’s usefulness in evaluating existing primers, we used it to assess previously published primers reported to be specific for the bacterial plant-pathogenic pathogenic taxa *Xanthomonas vesicatoria* and *Xanthomonas vasicola* pv. *vasculorum*. The previously published *Xanthomonas vesicatoria-*specific primer pair, Xv-gyrB-F (5ʹ-ATACGCGTTGGGCGAGCCT-3ʹ) and Xv-gyrB-R (5ʹ-CATCGCTGAAGATGGCCACG GCT-3ʹ)^27^, and the previously published *Xanthomonas vasicola* pv. *vasculorum*-specific primer pair, Xvv3-F (5’-CAAGCAGAGCATGGCAAAC-3’) and Xvv3-R (5’-CACGTAGAACCGGTCTTTGG-3’)^28^, were tested for specificity and sensitivity against a panel of more than 600 *Xanthomonas* assemblies (**Dataset S3**) using the testing-only mode of DiPPER2 with default parameter values. Metadata of neighbour contigs for which correct-sized amplicons were generated was retrieved using E-direct utilities (esearch, elink, efetch and xtract)^46^.

### Phylogenetic reconstruction and species delineation

PhaME was used to infer the phylogeny from whole-genome sequences ^30^. PhaME was run with BowTie2 as an aligner and FastTree for inferring approximately-maximum-likelihood phylogenetic trees ^44,48^. The number of bootstraps was set to 1000 and the threshold for SNP-calling was 0.6. Only assemblies with a CheckM completeness of over 90% and with less than 1% contamination were included in the tree (accession list in **Dataset S3**) ^29^. CheckM values were retrieved from NCBI where they were calculated on the Prokaryotic Genome Annotation Pipeline (PGAP) gene set with the *Pseudomonas* or *Xanthomonas* CheckM marker set, respectively ^29,54^. The tree was visualized using iTOL and annotated with the iTOL annotation editor ^31,49^.

To further investigate species delineation between genome assemblies (**Figure 3** and **Dataset S3**), the genome-to-genome distance calculator (GGDC) tool available on DSMZ’s Type (Strain) Genome Server (TYGS) was used to calculate digital DNA:DNA-hybridization (dDDH; d4 formula) values ^26^. ANI values were calculated for the same genomes using FastANI with default settings using the FastANI analysis protocol ^47,55^.

