## Supplemental Material for "DiPPER2 – a user-friendly pipeline for picking and evaluating taxon-specific PCR primers"

**This pdf includes:**

**Supplementary figures and tables:**
Figures S1 to S5 (page 2-6)
Table S1 and S2 (page 7-8)

**Supplementary datasets:**
Legends of Datasets S1 to S3 (page 9)

**References:**
References (page 10)

**Supplementary Figures:**

**A)**

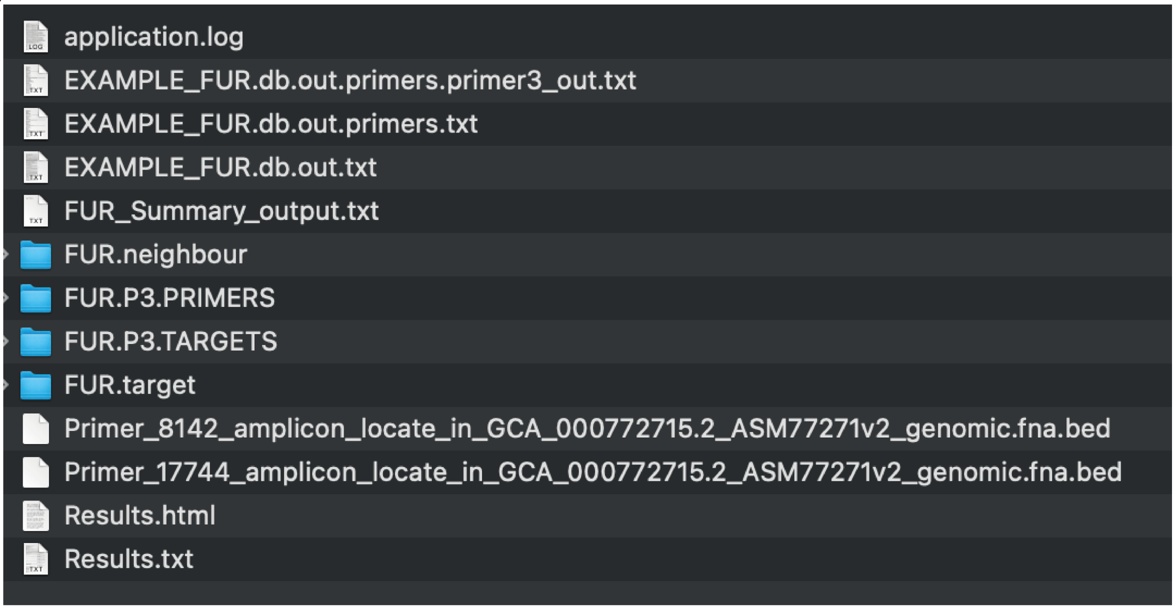


**B)**

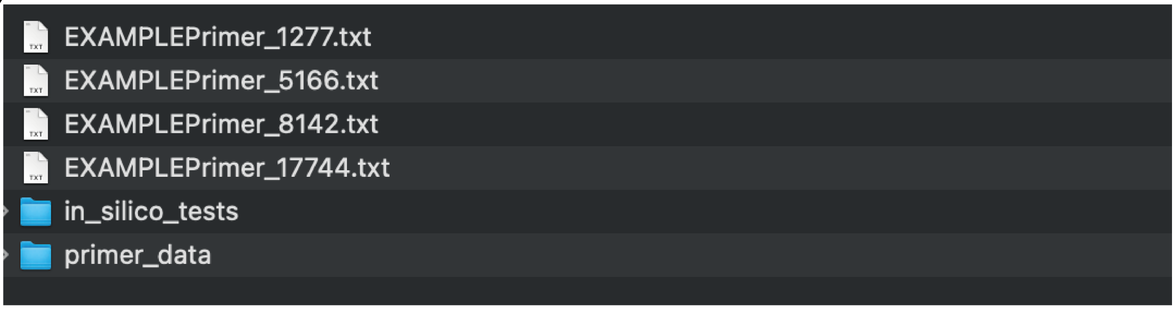


**Figure S1: `DiPPER2` results folder structure & files**

A) `DiPPER2` results folder. The logged run information is found in `application.log`. `Primer3` output (not reformatted) is found in the file with the suffix `*_FUR.db.out.primers.primer3_out.txt`. The `Primer3` input file with the unique genomic regions and `Primer3` parameterization is found in the file with the suffix `*_FUR.db.out.primers.txt`. The file with the suffix `*_FUR.db.out.txt` contains FURs output in fasta format which contains the unique genomic regions. `FUR.neighbour` is the folder with all neighbour assemblies. Similarly, `FUR.target` contains all target assemblies. `FUR.P3.PRIMERS` contains the primer pair fastas, *in silico* data and primer metadata. `FUR.P3.TARGETS` contains the fastas with the target regions of the respective primer pairs and the `blastx` results, if applicable. BED files are generated for primers for whom no `blastx` results were retrieved for their target sequence regions. The comprehensive results overview is found in Results.html and Results.txt

B) The `FUR.P3.PRIMERS` folder with its subfolders

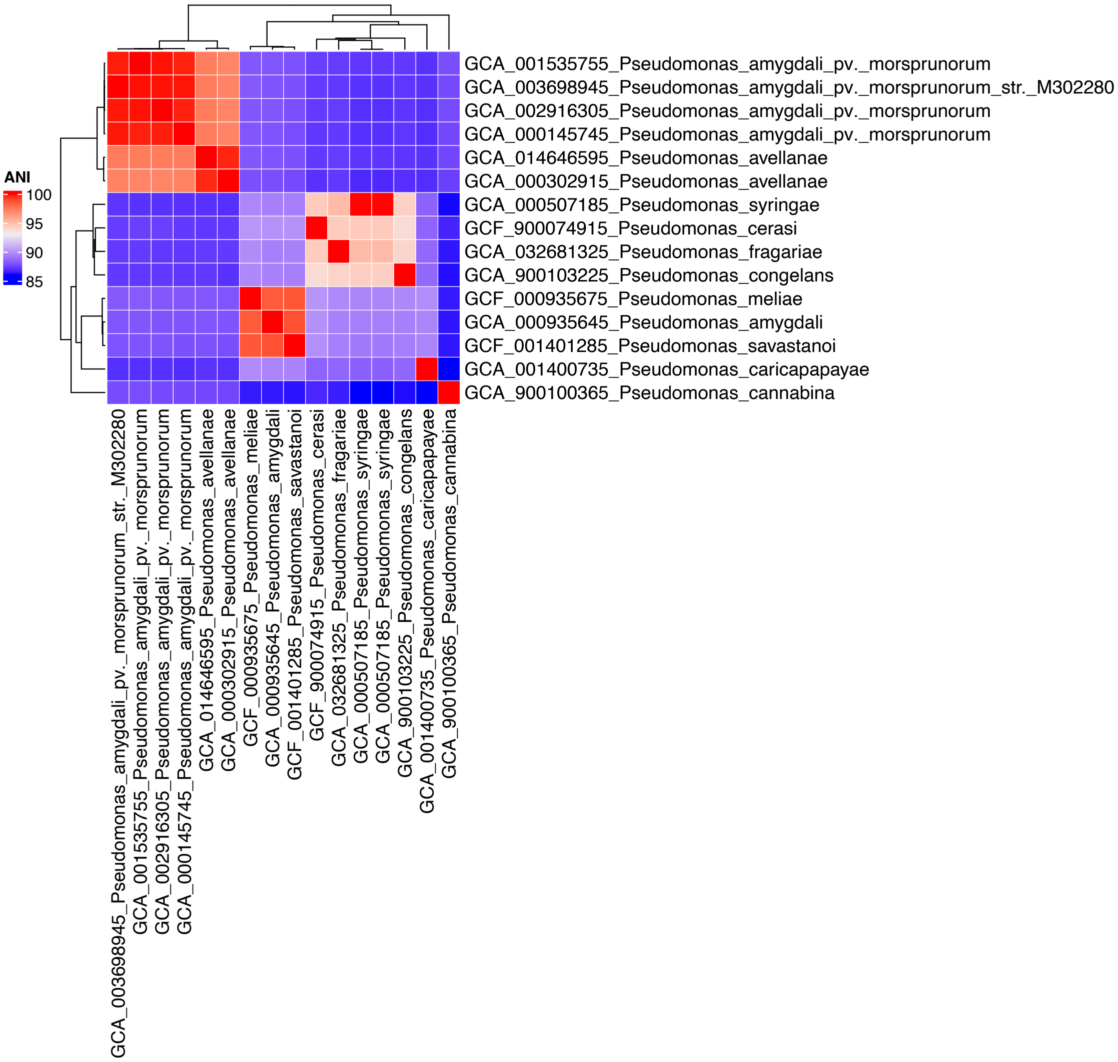


**Figure S2: FastANI[1] values represented in a heatmap.**

Lower values correspond with lower genetic homology and higher phylogenetic distance between strains. A threshold of 94-96% is widely accepted to delineate species boundaries[2–4], thus indicating that the four *Pseudomonas amygdali* pv. *morsprunorum* strains are closest related to *P. avellanae*. The strains in this analysis correspond to those in the TYGS analysis (**Dataset S2** and **3**).

### RESULTS FOR qPCR PRIMER 6689 in X_vasicola_qPCRPrimer_6689.txt

#### qPCR Primers

**Forward Primer:**
>Forward Primer 6689
CCATATCTGTGCGCCAAAGC

**Reverse Primer:**
>Reverse Primer 6689
GGAGTTGATGGGCGGTAACA

**Internal Probe:**
>Internal Probe 6689
GCGCGGCTCGGGTATCACGGT

#### Sensitivity and Specificity Testing

##### Sensitivity:

PASS OR FAIL:

```
PASSED
```

Number of assemblies the in silico PCR was run for:

```
6
```

Number of mismatches the sensitivity test failed for (m3 is tolerated):

```
m3
```

##### Specificity:

PASS OR FAIL:

```
PASSED
```

Number of assemblies the in silico PCR was run for:

```
NA
```

Number of mismatches the in silico PCR generated amplicons for:
( Note that if the specificity test has "passed", this means amplicons of incorrect lengths)

```
NA
```

#### BLASTX Target Testing

One or more files for *Target_6689.txt_blastx_1e-5.txt are empty. Blastx did not return result

#### Files & Folders

Primers are found in:

```
/Users/theresawacker/Desktop/workspace/Joanna_XaP_Xv_Xvv_qPCR/X_vasicola_qPCR/FUR.P3.PRIMERS
```

Target fastas and BLASTX results are found in:

```
/Users/theresawacker/Desktop/workspace/Joanna_XaP_Xv_Xvv_qPCR/X_vasicola_qPCR/FUR.P3.TARGETS
```

If generated, bed files are found in:

```
/Users/theresawacker/Desktop/workspace/Joanna_XaP_Xv_Xvv_qPCR/X_vasicola_qPCR
```

In silico PCR results are found in:

```
/Users/theresawacker/Desktop/workspace/Joanna_XaP_Xv_Xvv_qPCR/X_vasicola_qPCR/FUR.P3.PRIMERS/in_silico_tests
```

#### Sensitivity and Specificity Testing in Detail

##### Sensitivity:


```
{'Number of files (in silico PCR result files for different number of primer mismatches) tested:': 4,
 'Number of files that failed:': 1,
 'Number of files that passed:': 3,
 'X_vasicola_qPCRPrimer_6689.txt_seqkit_amplicon_against_target_m0.txt': {'Did the test pass?': 'passed',
                                                                            'Mismatches tested': 'm0',
                                                                            'Number of assemblies with correct size amplicon': 6,
                                                                            'Number of assemblies with wrong size amplicon': 0,
                                                                            'Number of assemblies, in silico PCR was performed on': 6},
 'X_vasicola_qPCRPrimer_6689.txt_seqkit_amplicon_against_target_m1.txt': {'Did the test pass?': 'passed',
                                                                            'Mismatches tested': 'm1',
                                                                            'Number of assemblies with correct size amplicon': 6,
                                                                            'Number of assemblies with wrong size amplicon': 0,
                                                                            'Number of assemblies, in silico PCR was performed on': 6},
 'X_vasicola_qPCRPrimer_6689.txt_seqkit_amplicon_against_target_m2.txt': {'Did the test pass?': 'passed',
                                                                            'Mismatches tested': 'm2',
                                                                            'Number of assemblies with correct size amplicon': 6,
                                                                            'Number of assemblies with wrong size amplicon': 0,
                                                                            'Number of assemblies, in silico PCR was performed on': 6},
 'X_vasicola_qPCRPrimer_6689.txt_seqkit_amplicon_against_target_m3.txt': {'Did the test pass?': 'failed',
                                                                            'Mismatches tested': 'm3',
                                                                            'Number of assemblies with correct size amplicon': 3,
                                                                            'Number of assemblies with wrong size amplicon': 3,
                                                                            'Number of assemblies, in silico PCR was performed on': 6}}
```


##### Specificity:


```
{'NA': {'Did the test pass?': 'NA',
        'Mismatches tested': 'NA',
        'Number of assemblies with correct size amplicon': 'NA',
        'Number of assemblies with wrong size amplicon': 'NA',
        'Number of assemblies, in silico PCR was performed on': 'NA'},
 'Number of files that failed:': 'NA',
 'Number of files that passed:': 'NA'}
```


**Figure S3: Example html report from DiPPER2 run**

### RESULTS FOR qPCR PRIMER 4209 in X_vasicola_qPCRPrimer_4209.txt

#### qPCR Primers

**Forward Primer:**
>Forward Primer 4209
TGCTTGATCGCTAAGGTGCA

**Reverse Primer:**
>Reverse Primer 4209
GCTGATCGACATTTCCGCAC

**Internal Probe:**
>Internal Probe 4209
GCGCTGCGCTTTGCCGTGGT

#### Sensitivity and Specificity Testing

##### Sensitivity:

PASS OR FAIL:

```
FAILED
```

Number of assemblies the in silico PCR was run for:

```
6
```

Number of mismatches the sensitivity test failed for (m3 is tolerated):

```
m3, m2
```

##### Specificity:

PASS OR FAIL:

```
PASSED
```

Number of assemblies the in silico PCR was run for:

```
NA
```

Number of mismatches the in silico PCR generated amplicons for:
( Note that if the specificity test has "passed", this means amplicons of incorrect lengths)

```
NA
```

#### BLASTX Target Testing

The primer 4209's target with the highest bitscore 115.0 & evalue 3.62e-31 has the accession WP_126922534.1 and codes for hypothetical protein [Xanthomonas vasicola]. Further information on the accession: https://www.ncbi.nlm.nih.gov/protein/WP_126922534.1/

#### Files & Folders

Primers are found in:

```
/Users/theresawacker/Desktop/workspace/Joanna_XaP_Xv_Xvv_qPCR/X_vasicola_qPCR/FUR.P3.PRIMERS
```

Target fastas and BLASTX results are found in:

```
/Users/theresawacker/Desktop/workspace/Joanna_XaP_Xv_Xvv_qPCR/X_vasicola_qPCR/FUR.P3.TARGETS
```

If generated, bed files are found in:

```
/Users/theresawacker/Desktop/workspace/Joanna_XaP_Xv_Xvv_qPCR/X_vasicola_qPCR
```

In silico PCR results are found in:

```
/Users/theresawacker/Desktop/workspace/Joanna_XaP_Xv_Xvv_qPCR/X_vasicola_qPCR/FUR.P3.PRIMERS/in_silico_tests
```

#### Sensitivity and Specificity Testing in Detail

##### Sensitivity:


```
{'Number of files (in silico PCR result files for different number of primer mismatches) tested:': 4,
 'Number of files that failed:': 2,
 'Number of files that passed:': 2,
 'X_vasicola_qPCRPrimer_4209.txt_seqkit_amplicon_against_target_m0.txt': {'Did the test pass?': 'passed',
                                                                          'Mismatches tested': 'm0',
                                                                          'Number of assemblies with correct size amplicon': 6,
                                                                          'Number of assemblies with wrong size amplicon': 0,
                                                                          'Number of assemblies, in silico PCR was performed on': 6},
 'X_vasicola_qPCRPrimer_4209.txt_seqkit_amplicon_against_target_m1.txt': {'Did the test pass?': 'passed',
                                                                          'Mismatches tested': 'm1',
                                                                          'Number of assemblies with correct size amplicon': 6,
                                                                          'Number of assemblies with wrong size amplicon': 0,
                                                                          'Number of assemblies, in silico PCR was performed on': 6},
 'X_vasicola_qPCRPrimer_4209.txt_seqkit_amplicon_against_target_m2.txt': {'Did the test pass?': 'failed',
                                                                          'Mismatches tested': 'm2',
                                                                          'Number of assemblies with correct size amplicon': 4,
                                                                          'Number of assemblies with wrong size amplicon': 2,
                                                                          'Number of assemblies, in silico PCR was performed on': 6},
 'X_vasicola_qPCRPrimer_4209.txt_seqkit_amplicon_against_target_m3.txt': {'Did the test pass?': 'failed',
                                                                          'Mismatches tested': 'm3',
                                                                          'Number of assemblies with correct size amplicon': 0,
                                                                          'Number of assemblies with wrong size amplicon': 9,
                                                                          'Number of assemblies, in silico PCR was performed on': 6}}
```


##### Specificity:


```
{'NA': {'Did the test pass?': 'NA',
        'Mismatches tested': 'NA',
        'Number of assemblies with correct size amplicon': 'NA',
        'Number of assemblies with wrong size amplicon': 'NA',
        'Number of assemblies, in silico PCR was performed on': 'NA'},
 'Number of files that failed:': 'NA',
 'Number of files that passed:': 'NA'}
```


**Figure S3: continued**

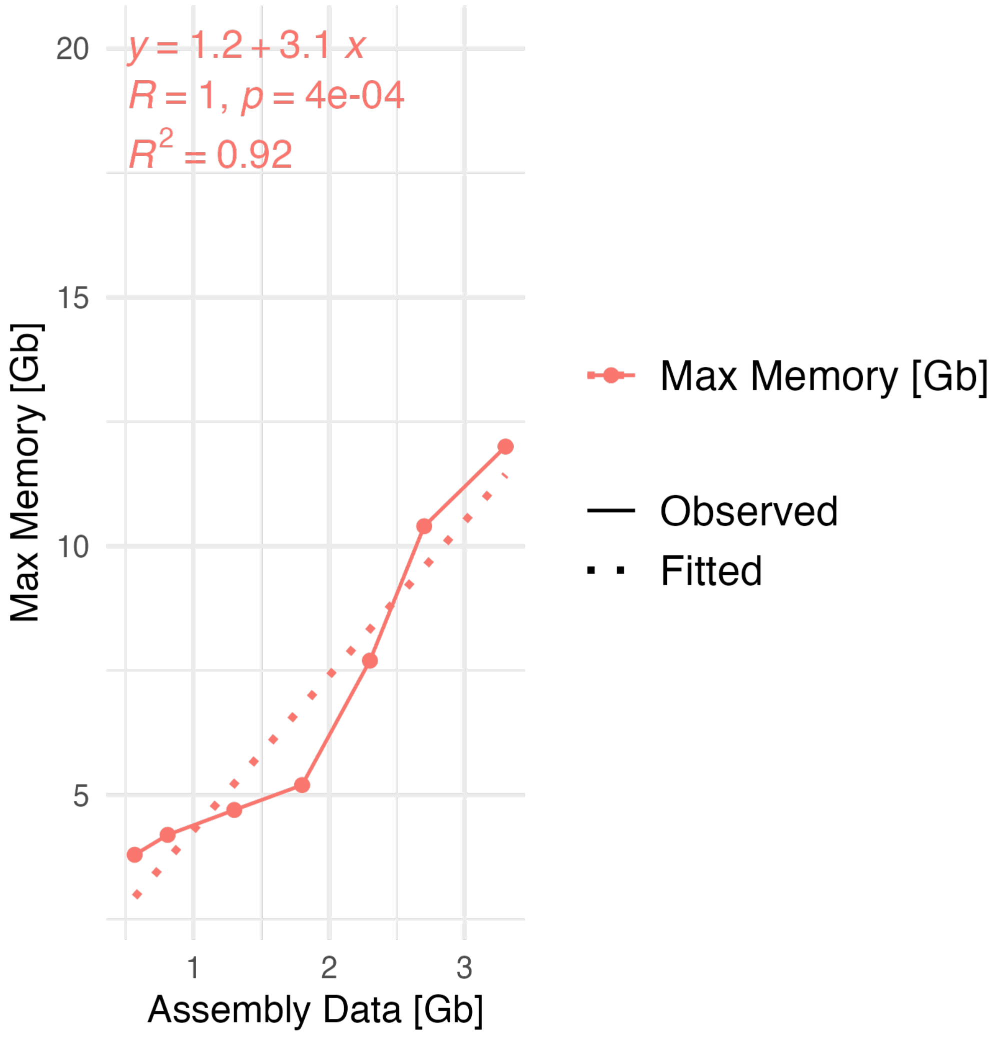


**Figure S4: Memory profile of DiPPER2 in full pipeline mode for up to 3.3 Gb of data**.
3.3 Gb of data are equivalent to 6 target assemblies and 660 neighbour assemblies. Max memory represents resident set size in Gigabytes (Gb). R= Spearman correlation coefficient; $R^2$=R-squared value/ coefficient of determination

#### Supplementary Tables:

##### Table S1: Metadata of the contigs (Acc.) of neighbours for which the *in silico* PCR generated correct-sized amplicons

| Acc: | Taxon: | ID: | BioSample: | SRA: | BioProject: | Strain: | Biomaterial provider: |
|---|---|---|---|---|---|---|---|
| CP066999.1 | *Xanthomonas campestris* pv. *campestris* | 17198903 | SAMN17198903 | SRS8608005 | PRJNA689092 | Xc_29_6 | USDA ARS- Ft. Detrick |
| CP066984.1 | *Xanthomonas campestris* pv. *campestris* | 17198915 | SAMN17198915 | SRS8607959 | PRJNA689092 | Xc_27_10 | USDA ARS- Ft. Detrick |
| CP066969.1 | *Xanthomonas campestris* | 17198925 | SAMN17198925 | SRS8607970 | PRJNA689092 | 42644 | Christine Smart- Cornell University |
| CP066985.1 | *Xanthomonas campestris* pv. *campestris* | 17198914 | SAMN17198914 | SRS8607958 | PRJNA689092 | Xc_25_4 | USDA ARS- Ft. Detrick |
| CP067000.1 | *Xanthomonas campestris* pv. *campestris* | 17198902 | SAMN17198902 | SRS8607994 | PRJNA689092 | Xc_29_5 | USDA ARS- Ft. Detrick |
| CP066986.1 | *Xanthomonas campestris* pv. *campestris* | 17198913 | SAMN17198913 | SRS8607957 | PRJNA689092 | Xc_15_1 | USDA ARS- Ft. Detrick |
| CP067001.1 | *Xanthomonas campestris* pv. *campestris* | 17198901 | SAMN17198901 | SRS8607983 | PRJNA689092 | Xc_40_2 | USDA ARS- Ft. Detrick |
| CP066983.1 | *Xanthomonas campestris* pv. *campestris* | 17198916 | SAMN17198916 | SRS8607960 | PRJNA689092 | Xc_27_6 | USDA ARS- Ft. Detrick |
| CP066982.1 | *Xanthomonas campestris* pv. *campestris* | 17198917 | SAMN17198917 | SRS8607962 | PRJNA689092 | Xc_27_9 | USDA ARS- Ft. Detrick |
| CP066978.1 | *Xanthomonas campestris* pv. *incanae* | 17198921 | SAMN17198921 | SRS8607966 | PRJNA689092 | 18048 | USDA ARS- Ft. Detrick |
| CM002636.1 | *Xanthomonas campestris* pv. *incanae* | 2645671 | SAMN02645671 | | PRJNA185977 | CFBP 2527R | LIPM, Toulouse, France |
| CP066928.1 | *Xanthomonas campestris* pv. *incanae* | 17198950 | SAMN17198950 | SRS8607998 | PRJNA689092 | CFBP1371 | |
| CP066981.1 | *Xanthomonas campestris* pv. *campestris* | 17198918 | SAMN17198918 | SRS8607963 | PRJNA689092 | Xc_28_1 | USDA ARS- Ft. Detrick |

**Table S2:Resource consumption of DiPPER2 in full pipeline mode**

Data = Gigabytes (Gb) of sequence data; n(neigh.) = number of neighbour assemblies; n(targets) = number of target assemblies; wall-clock = wall-clock time in min; cumul. CPU = cumulative CPU time over all cores in min; max(rss) = maximum resident set size, which represents the amount of memory (RAM) that the process uses; parall. = parallelized version of DiPPER2

| Data [Gb] | n(neigh.) | n(targets) | Wall-clock [min] | Cumul. CPU [min] | max(rss) [Gb] | Wall-clock parall. [min] | Cumul. CPU parall. [min] | max(rss) [Gb] |
|---|---|---|---|---|---|---|---|---|
| 3.3 | 660 | 6 | 118 | 362 | 12 | 147 | 355 | 9.9 |
| 2.7 | 560 | 6 | 105 | 308 | 10.4 | 130 | 303 | 8.1 |
| 2.3 | 460 | 6 | 87 | 250 | 7.7 | NAN | NAN | NAN |
| 1.8 | 360 | 6 | 72 | 199 | 5.2 | NAN | NAN | NAN |
| 1.3 | 260 | 6 | 50 | 144 | 4.7 | NAN | NAN | NAN |
| 0.809 | 160 | 6 | 72 | 90 | 4.2 | NAN | NAN | NAN |
| 0.566 | 110 | 6 | 51 | 64 | 3.8 | NAN | NAN | NAN |

**Dataset Legends:**

**Dataset S1. fastANI and dDDH values for *Pseudomonas*.**
**A)** fastANI[1] results with calculated average nucleotide identities (ANI) for the 4 mislabelled *Pseudomonas amygdali* pv. *morsprunorum* strains (highlighted in bold; **Dataset S3 B)** and **Figure 3**) and Type (strain) genome server (TYGS)[5] strains from type material.
**B)** fastANI[1] results from all *Pseudomonas* strains in the phylogenetic tree in **Figure 2**, which represent *Pseudomonas* strains on NCBI[6] that have a CheckM[7,8] completeness of ≥90% and ≤1% contamination (**Dataset S3 B)**). This includes the 4 mislabelled *Pseudomonas amygdali* pv. *morsprunorum* (highlighted in bold).
**C)** TYGS[5] pairwise dDDH values for the 4 mislabelled strains and the TYGS-chosen *Pseudomonas* strains from type material. dDDH = digital DNA:DNA hybridisation; C.I. = confidence interval; $d_0$, $d_3$, $d_4$ = GBDP (Genome BLAST Distance Phylogeny) formulas explained in **Dataset S2 Table 3.**

**Dataset S2. TYGS[5] results for the 4 *Pseudomonas amygdali* pv. *morsprunorum* strains**
Type (strain) genome server job result file which includes:
- Table 2: contains the result of the TYGS[5] species identification routine
- Table 3: containing the pairwise dDDH values
- Table 4: information about the strains
- Material, Methods and References

**Dataset S3. Strain accessions for *Xanthomonas* and *Pseudomonas***
A) *Xanthomonas* accessions of all strains used in the memory and runtime profiling full pipeline runs of `DiPPER2`. Target assemblies in this case were for the *X. vasicola* clade, the accessions are listed in column 2.
B) *Pseudomonas* accessions of all strains in the phylogenetic tree in **Figure 2**, which represent *Pseudomonas* strains on NCBI[6] that have a CheckM[7,8] completeness of ≥90% and ≤1% contamination. This includes the 4 mislabelled *Pseudomonas amygdali* pv. *morsprunorum*. Also included are the accessions of the *Pseudomonas* strains chosen by TYGS for the dDDH calculations.
C) Information about the strains used in the TYGS calculations.
