## Supplementary material for "DiPPER2 – a user-friendly pipeline for picking and evaluating taxon-specific PCR primers": Dataset S2

TYPE (STRAIN) GENOME SERVER

### JOB SUMMARY

PRINT DATE: 2024-10-02 14:54:29 +0200

JOB ID: 73325ce0-5cb9-423c-b501-78fba54f571e

RESULT PAGE: https://tygs.dsmz.de/user_results/show?guid=73325ce0-5cb9-423c-b501-78fba54f571e

#### Table 1: Phylogenies

**Publication-ready versions** of both the genome-scale GBDP tree and the 16S rRNA gene sequence tree can be customized and exported either in SVG (vector graphic) or PNG format from within the phylogeny viewers in your TYGS result page. For publications the **SVG format is recommended** because it is lossless, always keeps its high resolution and can also be easily converted to other popular formats such as PDF or EPS. Please follow the link provided above!

#### Table 2: Identification

The below list contains the result of the TYGS species identification routine.

Explanation of remarks that might occur in the below table:

**remark [R1]:** The TYGS type strain database is automatically updated on an almost daily basis. However, if a particular type strain genome is not available in the TYGS database, this can have several reasons which are detailed in the FAQ. You can request an extended 16S rRNA gene analysis via the 16S tree viewer found in your result page to detect **not yet genome-sequenced** type strains relevant for your study.

**remark [R2]:** > 70% dDDH value (formula $d_4$) and (almost) minimal dDDH values for gene-content formulae $d_0$ and $d_6$ indicate a potentially unreliable identification result and should thus be checked via the 16S rRNA gene sequence similarity. Such strong deviations can, in principle, be caused by sequence contamination.

**remark [R3]:** G+C content difference of > 1 % indicates a potentially unreliable identification result because within species G+C content varies no more than 1 %, if computed from genome sequences (PMID: 24505073).

| Strain | Conclusion | Identification result | Remark |
|---|---|---|---|
| **'Pseudomonas amygdali pv. morsprunorum str. M302280' (GCA_000145745.1)** | belongs to known species | *Pseudomonas avellanae* | |
| **'Pseudomonas amygdali pv. morsprunorum' (GCA_001535755.1)** | belongs to known species | *Pseudomonas avellanae* | |
| **'Pseudomonas amygdali pv. morsprunorum' (GCA_002916305.1)** | belongs to known species | *Pseudomonas avellanae* | |
| **'Pseudomonas amygdali pv. morsprunorum' (GCA_003698945.1)** | belongs to known species | *Pseudomonas avellanae* | |

#### Table 3: Pairwise comparisons of user genomes vs. type-strain genomes

The following table contains the pairwise dDDH values between your user genomes and the selected type-strain genomes. The dDDH values are provided along with their confidence intervals (C.I.) for the three different GBDP formulas:

- formula $d_0$ (a.k.a. GGDC formula 1): length of all HSPs divided by total genome length
- formula $d_4$ (a.k.a. GGDC formula 2): sum of all identities found in HSPs divided by overall HSP length
- formula $d_6$ (a.k.a. GGDC formula 3): sum of all identities found in HSPs divided by total genome length

**Note:** Formula $d_4$ is independent of genome length and is thus robust against the use of incomplete draft genomes. For other reasons for preferring formula $d_4$, see the FAQ.

| Query | Subject | $d_0$ | C.I. $d_0$ | $d_4$ | C.I. $d_4$ | $d_6$ | C.I. $d_6$ | Diff. G+C Percent |
|---|---|---|---|---|---|---|---|---|
| **'Pseudomonas amygdali pv. morsprunorum str. M302280' (GCA_000145745.1)** | **'Pseudomonas amygdali pv. morsprunorum' (GCA_003698945.1)** | 99.9 | [99.8 - 100.0] | 99.8 | [99.6 - 99.9] | 100.0 | [99.9 - 100.0] | 0.0 |
| **'Pseudomonas amygdali pv. morsprunorum' (GCA_001535755.1)** | **'Pseudomonas amygdali pv. morsprunorum' (GCA_002916305.1)** | 98.3 | [97.2 - 99.0] | 99.6 | [99.3 - 99.8] | 99.2 | [98.6 - 99.5] | 0.02 |
| **'Pseudomonas amygdali pv. morsprunorum str. M302280' (GCA_000145745.1)** | **'Pseudomonas amygdali pv. morsprunorum' (GCA_002916305.1)** | 98.0 | [96.7 - 98.8] | 99.4 | [99.1 - 99.6] | 99.0 | [98.4 - 99.4] | 0.21 |
| **'Pseudomonas amygdali pv. morsprunorum str. M302280' (GCA_000145745.1)** | **'Pseudomonas amygdali pv. morsprunorum' (GCA_001535755.1)** | 97.6 | [96.1 - 98.5] | 99.3 | [98.9 - 99.6] | 98.7 | [98.0 - 99.2] | 0.19 |
| **'Pseudomonas amygdali pv. morsprunorum' (GCA_002916305.1)** | **'Pseudomonas amygdali pv. morsprunorum' (GCA_003698945.1)** | 98.0 | [96.6 - 98.8] | 99.3 | [98.9 - 99.6] | 99.0 | [98.3 - 99.4] | 0.21 |
| **'Pseudomonas amygdali pv. morsprunorum' (GCA_001535755.1)** | **'Pseudomonas amygdali pv. morsprunorum' (GCA_003698945.1)** | 97.5 | [96.0 - 98.5] | 99.2 | [98.8 - 99.5] | 98.7 | [97.9 - 99.2] | 0.19 |
| **'Pseudomonas amygdali pv. morsprunorum str. M302280' (GCA_000145745.1)** | *Pseudomonas avellanae* BPIC 631 | 66.9 | [63.0 - 70.5] | 80.4 | [77.5 - 83.1] | 71.4 | [67.9 - 74.6] | 0.08 |
| **'Pseudomonas amygdali pv. morsprunorum' (GCA_003698945.1)** | *Pseudomonas avellanae* BPIC 631 | 67.1 | [63.2 - 70.7] | 80.1 | [77.2 - 82.7] | 71.5 | [68.0 - 74.7] | 0.08 |
| **'Pseudomonas amygdali pv. morsprunorum str. M302280' (GCA_000145745.1)** | *Pseudomonas avellanae* JCM 11937 | 77.4 | [73.5 - 81.0] | 79.8 | [76.9 - 82.5] | 80.7 | [77.3 - 83.7] | 0.07 |
| **'Pseudomonas amygdali pv. morsprunorum' (GCA_002916305.1)** | *Pseudomonas avellanae* BPIC 631 | 65.6 | [61.8 - 69.2] | 79.6 | [76.6 - 82.2] | 70.0 | [66.6 - 73.3] | 0.29 |
| **'Pseudomonas amygdali pv. morsprunorum' (GCA_001535755.1)** | *Pseudomonas avellanae* BPIC 631 | 66.5 | [62.7 - 70.2] | 79.2 | [76.3 - 81.9] | 70.8 | [67.4 - 74.1] | 0.27 |
| **'Pseudomonas amygdali pv. morsprunorum' (GCA_003698945.1)** | *Pseudomonas avellanae* JCM 11937 | 77.9 | [73.9 - 81.4] | 78.9 | [76.0 - 81.6] | 80.9 | [77.6 - 83.9] | 0.06 |
| **'Pseudomonas amygdali pv. morsprunorum' (GCA_002916305.1)** | *Pseudomonas avellanae* JCM 11937 | 77.3 | [73.3 - 80.8] | 78.4 | [75.5 - 81.1] | 80.3 | [76.9 - 83.3] | 0.15 |
| **'Pseudomonas amygdali pv. morsprunorum' (GCA_001535755.1)** | *Pseudomonas avellanae* JCM 11937 | 78.1 | [74.1 - 81.6] | 78.1 | [75.2 - 80.8] | 80.9 | [77.6 - 83.9] | 0.13 |
| **'Pseudomonas amygdali pv. morsprunorum' (GCA_002916305.1)** | *Pseudomonas savastanoi* ICMP 4352 | 65.1 | [61.3 - 68.7] | 34.1 | [31.6 - 36.6] | 57.0 | [53.9 - 60.2] | 0.42 |

| Query | Subject | $d_0$ | C.I. $d_0$ | $d_4$ | C.I. $d_4$ | $d_6$ | C.I. $d_6$ | Diff. G+C Percent |
|---|---|---|---|---|---|---|---|---|
| **'Pseudomonas amygdali pv. morsprunorum' (GCA_001535755.1)** | *Pseudomonas savastanoi* ICMP 4352 | 64.6 | [60.8 - 68.2] | 34.0 | [31.5 - 36.5] | 56.7 | [53.5 - 59.8] | 0.45 |
| **'Pseudomonas amygdali pv. morsprunorum' (GCA_001535755.1)** | *Pseudomonas amygdali* CFBP 3205 | 64.7 | [60.9 - 68.3] | 34.0 | [31.6 - 36.5] | 56.7 | [53.6 - 59.9] | 0.15 |
| **'Pseudomonas amygdali pv. morsprunorum' (GCA_002916305.1)** | *Pseudomonas amygdali* CFBP 3205 | 64.7 | [60.9 - 68.3] | 34.0 | [31.6 - 36.5] | 56.7 | [53.5 - 59.8] | 0.13 |
| **'Pseudomonas amygdali pv. morsprunorum str. M302280' (GCA_000145745.1)** | *Pseudomonas amygdali* CFBP 3205 | 63.6 | [59.8 - 67.2] | 33.9 | [31.5 - 36.4] | 55.9 | [52.7 - 59.0] | 0.34 |
| **'Pseudomonas amygdali pv. morsprunorum' (GCA_001535755.1)** | *Pseudomonas meliae* CFBP 3225 | 60.9 | [57.2 - 64.5] | 33.9 | [31.5 - 36.4] | 53.9 | [50.8 - 57.0] | 0.04 |
| **'Pseudomonas amygdali pv. morsprunorum' (GCA_002916305.1)** | *Pseudomonas meliae* CFBP 3225 | 60.5 | [56.8 - 64.1] | 33.8 | [31.4 - 36.3] | 53.6 | [50.4 - 56.7] | 0.02 |
| **'Pseudomonas amygdali pv. morsprunorum str. M302280' (GCA_000145745.1)** | *Pseudomonas savastanoi* ICMP 4352 | 62.9 | [59.1 - 66.5] | 33.8 | [31.4 - 36.3] | 55.3 | [52.1 - 58.4] | 0.64 |
| **'Pseudomonas amygdali pv. morsprunorum' (GCA_003698945.1)** | *Pseudomonas amygdali* CFBP 3205 | 65.2 | [61.4 - 68.8] | 33.8 | [31.3 - 36.3] | 57.0 | [53.8 - 60.1] | 0.34 |
| **'Pseudomonas amygdali pv. morsprunorum str. M302280' (GCA_000145745.1)** | *Pseudomonas meliae* CFBP 3225 | 60.7 | [57.0 - 64.3] | 33.8 | [31.4 - 36.3] | 53.7 | [50.6 - 56.8] | 0.23 |
| **'Pseudomonas amygdali pv. morsprunorum' (GCA_001535755.1)** | *Pseudomonas cannabina* ICMP 2823 | 58.5 | [54.9 - 62.1] | 33.7 | [31.3 - 36.2] | 52.1 | [49.0 - 55.1] | 0.02 |
| **'Pseudomonas amygdali pv. morsprunorum' (GCA_003698945.1)** | *Pseudomonas meliae* CFBP 3225 | 62.0 | [58.3 - 65.6] | 33.7 | [31.3 - 36.2] | 54.6 | [51.5 - 57.7] | 0.23 |
| **'Pseudomonas amygdali pv. morsprunorum str. M302280' (GCA_000145745.1)** | *Pseudomonas cannabina* ICMP 2823 | 57.0 | [53.5 - 60.5] | 33.7 | [31.3 - 36.2] | 50.9 | [47.9 - 54.0] | 0.18 |
| **'Pseudomonas amygdali pv. morsprunorum' (GCA_002916305.1)** | *Pseudomonas cannabina* ICMP 2823 | 58.5 | [54.9 - 62.1] | 33.6 | [31.2 - 36.2] | 52.0 | [48.9 - 55.1] | 0.04 |
| **'Pseudomonas amygdali pv. morsprunorum' (GCA_003698945.1)** | *Pseudomonas savastanoi* ICMP 4352 | 64.4 | [60.6 - 68.0] | 33.6 | [31.2 - 36.2] | 56.3 | [53.2 - 59.5] | 0.64 |
| **'Pseudomonas amygdali pv. morsprunorum' (GCA_003698945.1)** | *Pseudomonas cannabina* ICMP 2823 | 58.1 | [54.5 - 61.6] | 33.5 | [31.1 - 36.0] | 51.7 | [48.6 - 54.7] | 0.17 |
| **'Pseudomonas amygdali pv. morsprunorum' (GCA_002916305.1)** | *Pseudomonas cerasi* 58 | 61.5 | [57.8 - 65.1] | 33.0 | [30.5 - 35.5] | 53.9 | [50.8 - 57.0] | 0.36 |
| **'Pseudomonas amygdali pv. morsprunorum' (GCA_001535755.1)** | *Pseudomonas cerasi* 58 | 61.1 | [57.4 - 64.7] | 32.9 | [30.5 - 35.4] | 53.6 | [50.5 - 56.7] | 0.34 |
| **'Pseudomonas amygdali pv. morsprunorum str. M302280' (GCA_000145745.1)** | *Pseudomonas cerasi* 58 | 59.2 | [55.5 - 62.7] | 32.7 | [30.3 - 35.3] | 52.1 | [49.0 - 55.1] | 0.15 |
| **'Pseudomonas amygdali pv. morsprunorum' (GCA_003698945.1)** | *Pseudomonas cerasi* 58 | 61.2 | [57.5 - 64.7] | 32.5 | [30.1 - 35.0] | 53.4 | [50.3 - 56.5] | 0.15 |

| Query | Subject | $d_0$ | C.I. $d_0$ | $d_4$ | C.I. $d_4$ | $d_6$ | C.I. $d_6$ | Diff. G+C Percent |
|---|---|---|---|---|---|---|---|---|
| **'Pseudomonas amygdali pv. morsprunorum str. M302280' (GCA_000145745.1)** | *Pseudomonas fragariae* 17 | 56.6 | [53.0 - 60.1] | 32.1 | [29.7 - 34.6] | 49.9 | [46.9 - 53.0] | 0.59 |
| **'Pseudomonas amygdali pv. morsprunorum str. M302280' (GCA_000145745.1)** | *Pseudomonas congelans* DSM 14939 | 60.6 | [56.9 - 64.2] | 32.0 | [29.6 - 34.5] | 52.7 | [49.6 - 55.8] | 0.69 |
| **'Pseudomonas amygdali pv. morsprunorum' (GCA_001535755.1)** | *Pseudomonas fragariae* 17 | 57.4 | [53.8 - 60.9] | 31.9 | [29.5 - 34.4] | 50.3 | [47.3 - 53.4] | 0.78 |
| **'Pseudomonas amygdali pv. morsprunorum' (GCA_002916305.1)** | *Pseudomonas congelans* DSM 14939 | 59.4 | [55.8 - 63.0] | 31.8 | [29.4 - 34.3] | 51.8 | [48.7 - 54.9] | 0.91 |
| **'Pseudomonas amygdali pv. morsprunorum' (GCA_001535755.1)** | *Pseudomonas congelans* DSM 14939 | 59.5 | [55.9 - 63.0] | 31.8 | [29.4 - 34.3] | 51.8 | [48.7 - 54.9] | 0.89 |
| **'Pseudomonas amygdali pv. morsprunorum' (GCA_003698945.1)** | *Pseudomonas congelans* DSM 14939 | 62.4 | [58.7 - 66.0] | 31.8 | [29.4 - 34.3] | 53.9 | [50.8 - 57.0] | 0.7 |
| **'Pseudomonas amygdali pv. morsprunorum str. M302280' (GCA_000145745.1)** | *Pseudomonas syringae* KCTC 12500 | 56.3 | [52.7 - 59.8] | 31.8 | [29.3 - 34.3] | 49.5 | [46.4 - 52.5] | 0.29 |
| **'Pseudomonas amygdali pv. morsprunorum' (GCA_003698945.1)** | *Pseudomonas fragariae* 17 | 58.6 | [55.0 - 62.1] | 31.8 | [29.4 - 34.4] | 51.2 | [48.1 - 54.3] | 0.6 |
| **'Pseudomonas amygdali pv. morsprunorum' (GCA_002916305.1)** | *Pseudomonas fragariae* 17 | 56.3 | [52.7 - 59.8] | 31.8 | [29.4 - 34.3] | 49.6 | [46.5 - 52.6] | 0.81 |
| **'Pseudomonas amygdali pv. morsprunorum str. M302280' (GCA_000145745.1)** | *Pseudomonas caricapapayae* ICMP 2855 | 60.8 | [57.1 - 64.4] | 31.7 | [29.3 - 34.3] | 52.7 | [49.6 - 55.8] | 0.31 |
| **'Pseudomonas amygdali pv. morsprunorum' (GCA_002916305.1)** | *Pseudomonas caricapapayae* ICMP 2855 | 61.2 | [57.5 - 64.8] | 31.7 | [29.3 - 34.2] | 53.0 | [49.9 - 56.1] | 0.1 |
| **'Pseudomonas amygdali pv. morsprunorum' (GCA_001535755.1)** | *Pseudomonas syringae* KCTC 12500 | 56.1 | [52.6 - 59.6] | 31.6 | [29.2 - 34.1] | 49.3 | [46.3 - 52.4] | 0.49 |
| **'Pseudomonas amygdali pv. morsprunorum' (GCA_003698945.1)** | *Pseudomonas syringae* KCTC 12500 | 58.5 | [54.9 - 62.0] | 31.6 | [29.2 - 34.1] | 51.0 | [47.9 - 54.1] | 0.3 |
| **'Pseudomonas amygdali pv. morsprunorum' (GCA_002916305.1)** | *Pseudomonas syringae* KCTC 12500 | 55.6 | [52.1 - 59.1] | 31.6 | [29.2 - 34.1] | 48.9 | [45.9 - 52.0] | 0.51 |
| **'Pseudomonas amygdali pv. morsprunorum' (GCA_001535755.1)** | *Pseudomonas caricapapayae* ICMP 2855 | 61.7 | [58.0 - 65.3] | 31.6 | [29.2 - 34.1] | 53.3 | [50.2 - 56.4] | 0.12 |
| **'Pseudomonas amygdali pv. morsprunorum' (GCA_003698945.1)** | *Pseudomonas caricapapayae* ICMP 2855 | 63.2 | [59.5 - 66.8] | 31.5 | [29.1 - 34.1] | 54.4 | [51.2 - 57.5] | 0.31 |

#### Table 4: Strains in your dataset

Joint dataset of automatically determined closest type strains (if this mode was chosen), manually selected type strains (if selected accordingly) and the provided user strains, if provided (marked in yellow).

| Strain | Authority | Other deposits | Synonyms | Base pairs | Percent G+C | No. proteins | Goldstamp | Bioproject accession | Biosample accession | Assembly accession | IMG OID |
|---|---|---|---|---|---|---|---|---|---|---|---|
| *Pseudomonas cerasi* 58 | Kałuzna et al. 2017 | CFBP 8305; LMG 28609 | *Pseudomonas cerasi* | 6345 254 | 58.8 | 5770 | | PRJNA224116 | SAMEA3894894 | GCF_900074915 | |
| *Pseudomonas avellanae* BPIC 631 | Janse et al. 1997 | CIP 105176; NCPPB 3487; DSM 11809; JCM 11937; BPIC 631; F11 | *Pseudomonas avellanae* | 5847 420 | 58.7 | 4713 | Gp0020557 | PRJNA84293 | SAMN02471966 | GCA_000302915 | 2531839710 |
| *Pseudomonas amygdali* CFBP 3205 | Psallidas and Panagopoulos 1975 | CIP 106734; NCPPB 2607; ICMP 3918; ATCC 33614; CCUG 32770; DSM 7298; LMG 13184; LMG 2123 | *Pseudomonas amygdali* | 5719 124 | 58.3 | 5288 | Gp0120625 | PRJNA274666 | SAMN03328995 | GCA_000935645 | |
| *Pseudomonas syringae* KCTC 12500 | van Hall 1902 | CFBP 1392; CIP 106698; NCPPB 281; NRRL B-1631; ICMP 3023; ATCC 19310; CCUG 14279; DSM 10604; LMG 1247; NCAIM B.01398 | *Pseudomonas syringae*; *Pseudomonas syringae* subsp. *syringae* | 6150 048 | 58.9 | 5082 | Gp0070928 | PRJNA227265 | SAMN02404616 | GCA_000507185 | 2571042114 |

| Strain | Authority | Other deposits | Synonyms | Base pairs | Percent G+C | No. proteins | Goldstamp | Bioproject accession | Biosample accession | Assembly accession | IMG OID |
|---|---|---|---|---|---|---|---|---|---|---|---|
| *Pseudomonas avellanae* JCM 11937 | Janse et al. 1997 | CIP 105176; NCPPB 3487; DSM 11809; JCM 11937; BPIC 631; F11 | *Pseudomonas avellanae* | 5944 692 | 58.6 | 5482 | | PRJDB10510 | SAMD00245224 | GCA_014646595 | |
| *Pseudomonas cannabina* ICMP 2823 | (ex Šutič and Dowson 1959) Gardan et al. 1999 emend. Bull et al. 2010 | CFBP 2341; CIP 106140; NCPPB 1437; DSM 16822; LMG 5096 | *Pseudomonas cannabina* | 6388 314 | 58.5 | 5635 | Gp0127163 | PRJEB15992 | SAMN05216597 | GCA_900100365 | |
| *Pseudomonas meliae* CFBP 3225 | Ogimi 1981 | NCPPB 3033; ICMP 6289; ATCC 33050; DSM 6759; MAFF 301463; LMG 2220; No. 2; Ogimi 2 | *Pseudomonas meliae* | 5143 623 | 58.4 | 4803 | Gp0120629 | PRJNA224116 | SAMN03328996 | GCF_000935675 | |
| *Pseudomonas savastanoi* ICMP 4352 | (Janse 1982) Gardan et al. 1992 | CFBP 1670; CIP 103721; NCPPB 639; ATCC 13522; DSM 19341; DSM 50298; LMG 2209 | *Pseudomonas savastanoi*; *Pseudomonas syringae* subsp. *savastanoi* | 6022 960 | 58.0 | 5578 | Gp0146403 | PRJNA224116 | SAMN03976268 | GCF_001401285 | |
| *Pseudomonas caricapapayae* ICMP 2855 | Robbs 1956 | CFBP 3204; CIP 106736; NCPPB 1873; ATCC 33615; CCUG 32775; DSM 21109; LMG 2152 | *Pseudomonas caricapapayae* | 6257 512 | 58.3 | 5542 | Gp0146395 | PRJNA292453 | SAMN03976290 | GCA_001400735 | |

| Strain | Authority | Other deposits | Synonyms | Base pairs | Percent G+C | No. proteins | Goldstamp | Bioproject accession | Biosample accession | Assembly accession | IMG OID |
|---|---|---|---|---|---|---|---|---|---|---|---|
| *Pseudomonas congelans* DSM 14939 | Behrendt et al. 2003 | LMG 21466; P 538/23 | *Pseudomonas congelans* | 5731 984 | 59.3 | 4980 | Gp0127158 | PRJEB16279 | SAMN05216596 | GCA_900103225 | |
| *Pseudomonas fragariae* 17 | Marin et al. 2024 | DSM 113340; LMG 32456 | *Pseudomonas fragariae* | 6103 421 | 59.2 | 5175 | | PRJNA1021223 | SAMN37546059 | GCA_032681325 | |
| 'Pseudomonas amygdali pv. morsprunorum str. M302280' (GCA_00014574 5.1) | | | | 6039 297 | 58.6 | 5838 | | | | | |
| 'Pseudomonas amygdali pv. morsprunorum' (GCA_00153575 5.1) | | | | 6470 123 | 58.4 | 6089 | | | | | |
| 'Pseudomonas amygdali pv. morsprunorum' (GCA_00291630 5.1) | | | | 6498 711 | 58.4 | 6101 | | | | | |
| 'Pseudomonas amygdali pv. morsprunorum' (GCA_00369894 5.1) | | | | 6077 867 | 58.6 | 5552 | | | | | |

#### Methods, Results and References

The genome sequence data were uploaded to the Type (Strain) Genome Server (TYGS), a free bioinformatics platform available under https://tygs.dsmz.de, for a whole genome-based taxonomic analysis [1]. The analysis also made use of recently introduced methodological updates and features [2]. Information on nomenclature, synonymy and associated taxonomic literature was provided by TYGS's sister database, the List of Prokaryotic names with Standing in Nomenclature (LPSN, available at https://lpsn.dsmz.de) [2]. The results were provided by the TYGS on 2024-10-02. The TYGS analysis was subdivided into the following steps:

##### Determination of closely related type strains

Determination of closest type strain genomes was done in two complementary ways: First, all user genomes were compared against all type strain genomes available in the TYGS database via the MASH algorithm, a fast approximation of intergenomic relatedness [3], and, the ten type strains with the smallest MASH distances chosen per user genome. Second, an additional set of ten closely related type strains was determined via the 16S rDNA gene sequences. These were extracted from the user genomes using RNAmmer [4] and each sequence was subsequently BLASTed [5] against the 16S rDNA gene sequence of each of the currently 21645 type strains available in the TYGS database. This was used as a proxy to find the best 50 matching type strains (according to the bitscore) for each user genome and to subsequently calculate precise distances using the Genome BLAST Distance Phylogeny approach (GBDP) under the algorithm 'coverage' and distance formula $d_5$ [6]. These distances were finally used to determine the 10 closest type strain genomes for each of the user genomes.

##### Pairwise comparison of genome sequences

For the phylogenomic inference, all pairwise comparisons among the set of genomes were conducted using GBDP and accurate intergenomic distances inferred under the algorithm 'trimming' and distance formula $d_5$ [6]. 100 distance replicates were calculated each. Digital DDH values and confidence intervals were calculated using the recommended settings of the GGDC 4.0 [2,6].

##### Phylogenetic inference

The resulting intergenomic distances were used to infer a balanced minimum evolution tree with branch support via FASTME 2.1.6.1 including SPR postprocessing [7]. Branch support was inferred from 100 pseudo-bootstrap replicates each. The trees were rooted at the midpoint [8] and visualized with PhyD3 [9].

##### Type-based species and subspecies clustering

The type-based species clustering using a 70% dDDH radius around each of the 11 type strains was done as previously described [1]. The resulting groups are shown in Table 1 and 4. Subspecies clustering was done using a 79% dDDH threshold as previously introduced [10].

#### Results

##### Type-based species and subspecies clustering

The resulting species and subspecies clusters are listed in Table 4, whereas the taxonomic identification of the query strains is found in Table 1. Briefly, the clustering yielded 8 species clusters and the provided query strains were assigned to 1 of these. Moreover, user strains were located in 1 of 8 subspecies clusters.

##### Figure caption SSU tree

**Figure 1.** Tree inferred with FastME 2.1.6.1 [7] from GBDP distances calculated from 16S rDNA gene sequences. The branch lengths are scaled in terms of GBDP distance formula $d_5$. The numbers above branches are GBDP pseudo-bootstrap support values > 60 % from 100 replications, with an average branch support of 58.4 %. The tree was rooted at the midpoint [8].

##### Figure caption genome tree

**Figure 2.** Tree inferred with FastME 2.1.6.1 [7] from GBDP distances calculated from genome sequences. The branch lengths are scaled in terms of GBDP distance formula $d_5$. The numbers above branches are GBDP pseudo-bootstrap support values > 60 % from 100 replications, with an average branch support of 84.4 %. The tree was rooted at the midpoint [8].
